# Local conformational plasticity underlies ligand recognition and proton coupling in MFS multidrug transporters

**DOI:** 10.64898/2026.05.25.726547

**Authors:** Kyo Coppieters ‘t Wallant, Thomas Tilmant, Romain Malempré, Maria Mamkaeva, Loïc Quinton, Cédric Govaerts, Chloé Martens

## Abstract

Polyspecific substrate recognition drives multidrug efflux and antibiotic resistance, yet its molecular basis remains unclear. Here, we use HDX-MS to compare ligand-dependent local dynamics in three multidrug efflux pumps: NorA, QacA and LmrP. In the apo state, all three display high flexibility in specific transmembrane helices, unlike homologous transporters with narrow substrate profiles. Substrate binding remodels these flexible regions, but in a transporter-specific manner, revealing divergent local adaptations within a conserved fold. In LmrP, protonation-mimicking mutation of a conserved acidic residue recapitulates the substrate-induced dynamic changes, supporting a model in which transmembrane helix flexibility couples protonation and substrate binding to the conformational changes required for transport. Together, our study identifies local plasticity beyond what is captured by static high-resolution structures, as an overlooked feature of polyspecific ligand recognition.

## Introduction

Active efflux of cytotoxic compounds constitutes a first-line defense strategy in bacteria and plays a central role in the emergence of antimicrobial resistance^1–4^. In Gram-positive organisms, multidrug efflux pumps belonging to the Major Facilitator Superfamily (MFS MDRs) represent the largest class of efflux systems^5^. These transporters recognize a remarkably broad spectrum of substrates, which are typically amphipathic and positively charged^6^. Chemically distinct antibiotics such as the macrolide erythromycin and tetracycline can be transported by the same protein, whereas other antibiotics, such as ampicillin, are not^7,8^. How the conserved MFS fold accommodates such substrate polyspecificity at the molecular level remains poorly understood^8^.

Recent advances based on deep mutational scanning of polyspecific transporters have systematically explored sequence-phenotype relationships and revealed that sequence encoded promiscuity extends to sites distant from residues involved in ligand recognition, pointing toward allosteric mechanism to enable polyspecific recognition^9–11^. However, while these screens highlight correlations between sequence and phenotype, they underscore a gap in our understanding of how allosteric communication is encoded within the protein. We postulate that this is encoded in protein local dynamics.

While X-ray crystallography and cryo-EM have yielded detailed structural frameworks for MFS transporters^5^, they often fail to fully capture local conformational plasticity, as structural heterogeneity may be compressed into consensus models or lost through reduced local resolution in flexible regions^12,13^. These so-called structural dynamics, including segmental flexibility, order-to-disorder transitions, and local fluctuations, must be considered to understand protein molecular mechanisms. Hydrogen–deuterium exchange coupled to mass spectrometry (HDX-MS) is particularly well suited to probing the dynamics of transport proteins^14–17^. HDX-MS measures the rate of hydrogen–deuterium exchange via the resulting mass increase quantified by mass spectrometry^18^. Exchange of labile backbone amide protons depends on hydrogen-bond stability and accessibility to bulk deuterium^14^. Consequently, changes in HDX rates report on alterations in solvent exposure and/or interactions involving backbone amides, including local secondary structure elements^19^. Throughout this work, we use the term “*local structural dynamics*” to describe such transient, reversible rearrangements of secondary structure.

To understand how local structural dynamics shape the molecular mechanisms of MFS MDRs transporters, we mapped hydrogen–deuterium exchange in three well-characterized members of that family: QacA, NorA, and LmrP. NorA^20^ and QacA^21^ are MDR efflux pumps expressed by *Staphylococcus aureus*, the deadliest gram-positive pathogen listed as a high priority by the WHO for research regarding new therapies^22^. LmrP, found in *Lactococcus lactis*, is a model MDR efflux pump that has been extensively characterized both structurally and biochemically^23–25^. LmrP and NorA^26,27^ display the canonical MFS architecture, consisting of twelve transmembrane helices arranged into two pseudo-symmetric six-helix bundles^28,29^, while QacA^30^ displays an additional pair of transmembrane helices. In all cases, the N-lobe and C-lobe rotate around a central mostly hydrophobic cavity to allow alternate access from either side of the membrane^26,27,30,31^. Like other MFS MDR efflux pumps, they are proton-coupled and use the energy stored in the transmembrane proton gradient to power active efflux^8^. The coupling between drug binding and proton import relies on the modulation of the conformational landscape by specific conserved residues, whose protonation state modulates interactions between the N-lobe and the C-lobe, thus stabilizing specific states in transport cycle ^23,27^.

Here, we systematically map the local structural dynamics of QacA, NorA and LmrP and observe substantial H/D exchange within transmembrane α-helices, revealing an unexpected degree of intrinsic flexibility that is not apparent from high-resolution structures and has not been reported in previous H/D exchange studies of other MFS transporters^32–35^. We further show that this local flexibility is remodelled by substrate binding and protonation, identifying an overlooked layer of regulation of the conformational changes that underlie transport.

## Results

### High Local Flexibility of Transmembrane Helices Revealed by HDX-MS

We performed bottom-up HDX-MS measurements on QacA, NorA and LmrP, purified to homogeneity in detergent micelles (**Methods**). After isotopic labeling at different time points (ranging from 30s to 1h) the three proteins were proteolyzed with pepsin for fragment separation and MS analysis (**Methods**). Optimization of the HDX-MS workflow allowed a sequence coverage exceeding 75% for each protein (**Supplementary Fig. 1**). First, we measured the local HDX rate over time under apo conditions on the detergent-solubilized wild-type proteins for at least three biological replicates (**Supplementary Figs. 2, 3** and **4**). The analysis of the relative fractional uptake (R.F.U.) values mapped onto the protein sequence reveals the local structural dynamics profile of the three proteins (**Fig. 1**). Notably, for all three proteins, peptides belonging to regions depicted as transmembrane helices (TMHs) display deuterium exchange at rates similar to loop regions of the proteins.

**Figure 1:**
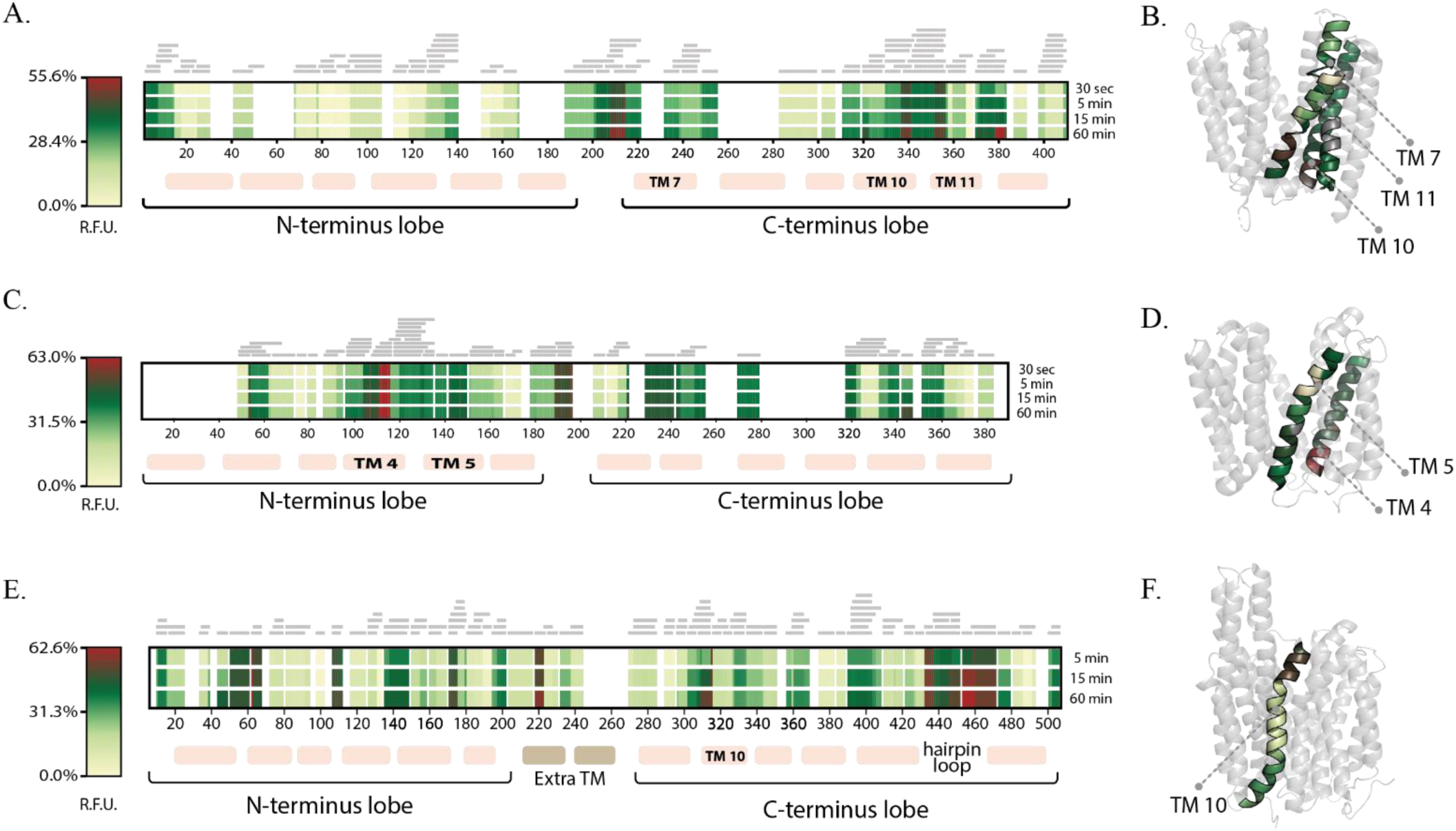
HDX-MS heatmaps of LmrP, NorA and QacA. Deuteration levels, fitted into heatmaps, of LmrP (A), NorA (C) and QacA (E) at the indicated time points. The transmembrane helices (TMHs) are respectively shown below the heatmaps with the TMHs containing highly exchanging peptides indicated. (B) The deuteration level after 60 min labelling is shown on the TMHs 7,10,11 of LmrP, identified as containing highly exchanging peptides. Data is plotted on the crystal structure (PDB : 61TZ). (D) The deuteration level after 60 min labelling is shown on the TMHs 4,5 of NorA, identified as containing highly exchanging peptides. Data are plotted on the cryo-EM structure (PDB : 7LO8). (F) The deuteration level after 60 min labelling is shown on the TMH 10 of QacA, identified as containing highly exchanging peptides. Data is plotted on the cryo-EM structure (PDB : 7Y58).

To validate the observed apparent transmembrane structural flexibility, we classified the peptides as “extramembranous” (when they included a loop or part of a loop) or “transmembranous” (when they were entirely contained within a TM helix) based on the three-dimensional structures of the proteins in both outward- and inward-facing conformations (**Methods**). This analysis resulted in curated datasets for each protein (**Supplementary Table 1, 2 and 3 and Supplementary Fig. 5**), for each biological replicate. We compared the exchange profiles of transmembrane peptides to the mean R.F.U. of unstructured and solvent-exposed loop regions, which served as a reference for maximal exchange. Peptides exceeding this threshold were classified as ‘**intrinsically dynamic’**, consistent with a local environment in which backbone amide protons are as accessible as those in non-helical regions. Because variability was substantial for each transporter, only peptides identified as dynamic in at least three biological replicates were classified as such, ensuring that these assignments reflected reproducible features rather than sample-specific variation.

The data shows that LmrP exhibits high exchange in transmembrane peptides of TMHs 7, 10, and 11 (**Fig. 1A, B, Supplementary. Fig.2 and Supplementary Table 1).** NorA presents high R.F.U. values across biological replicates in TMHs 4 and 5 (**Fig. 1C, D Supplementary Fig.3 and Suppl. Table 2**), while QacA consistently shows high values of HDX in transmembrane peptides of TM10 (**Fig. 1E, F, Supplementary Fig.4 and Supplementary Table 3**). Notably, specific segments within the transmembrane core, exemplified by peptides 231-238 (TMH7) and 356-365 (TMH11) in LmrP, and peptide 108-115 (TMH4) in NorA displayed a fast exchange regime. These regions reached high R.F.U. levels as early as the 30-second timepoint (**Fig. 2B, D, F**). Such rapid kinetics indicate frequent transient unfolding events, even within segments depicted as membrane-embedded α-helices. Overall, the analysis of deuterium uptake profiles of the apo proteins in detergent micelles clearly indicates that transmembrane α-helices exhibit disruptions in their hydrogen-bonding network.

**Figure 2.**
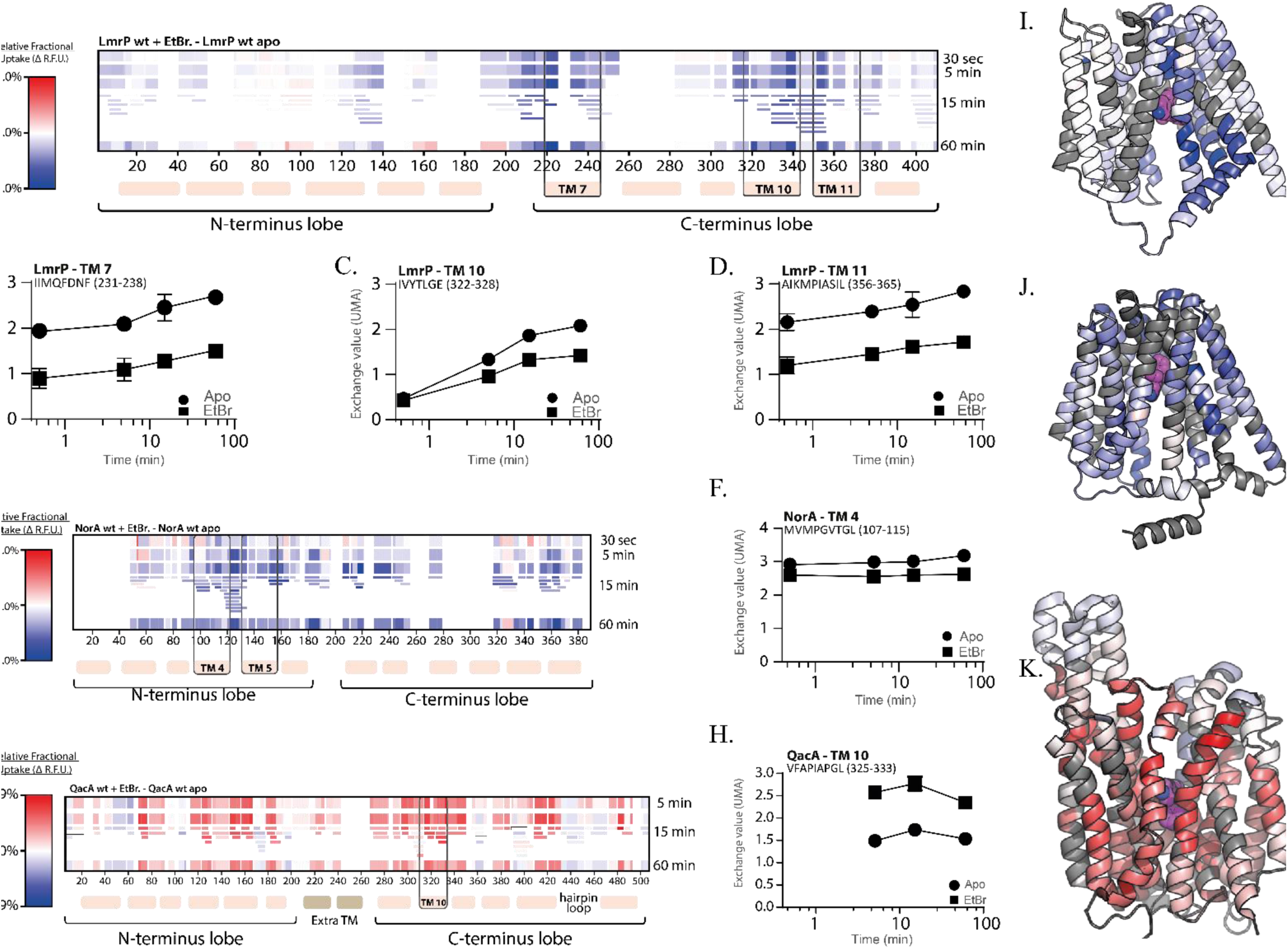
Ethidium bromide reshapes local exchange patterns across multidrug MFS transporters. (A) ΔR.F.U. heatmap comparing LmrP WT in the presence of ethidium bromide (EtBr) with apo LmrP WT. Values are shown on a blue–white–red scale, with blue indicating reduced exchange and red indicating increased exchange upon EtBr addition. (B–D, Deuterium uptake plots for representative peptides from TMH7, TMH10 and TMH11 of LmrP, comparing apo protein (circles) and EtBr-bound protein (squares). (E) ΔR.F.U. heatmap comparing NorA WT + EtBr with apo NorA WT. (F) Corresponding uptake plot for a representative peptide from TMH4 of NorA. (G) ΔR.F.U. heatmap comparing QacA WT + EtBr with apo QacA WT. (H) Corresponding uptake plot for a representative peptide from TMH10 of QacA. The peptides shown in the uptake plots correspond to regions previously identified as intrinsically flexible. (I–K) mapping of 15-min ΔR.F.U. values onto structural models of ligand-bound LmrP and NorA generated with Chai-1^41^, and onto the crystal structure of QacA bound to ethidium (PDB: 29PX).

**Figure 3.**
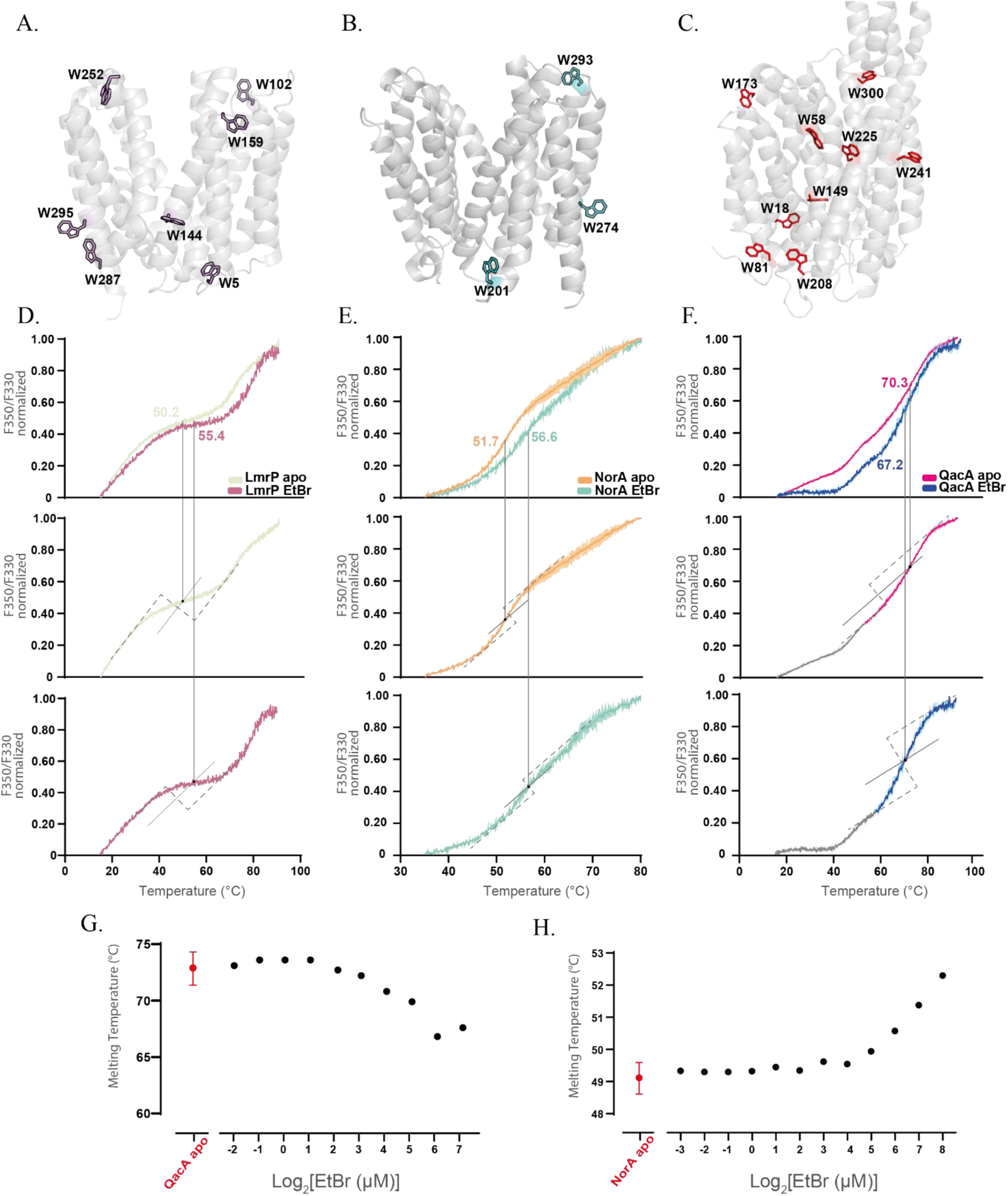
Ethidium binding differentially affects the thermal unfolding profiles of MFS multidrug transporters. (A–C). Structural mapping of intrinsic tryptophan residues in LmrP, NorA and QacA, respectively. Tryptophan residues are shown as sticks and coloured by transporter. (D–F). nanoDSF thermal unfolding profiles of LmrP, NorA and QacA in the absence or presence of 125 µM ethidium bromide (EtBr), monitored as the normalized F350/F330 fluorescence ratio. Curves represent the mean of two technical replicates, with shaded areas indicating the deviation between replicates. Apparent transition temperatures were extracted using a tangent-extrapolation method, as described in the Methods. Representative tangent constructions are shown below the corresponding unfolding curves. (G). EtBr concentration dependence of the second apparent transition temperature, Tapp2, for QacA. (H). EtBr concentration dependence of the apparent transition temperature, Tapp, for NorA. Titrations were performed over EtBr concentration ranges of 0.25–128 µM for QacA and 0.125–256 µM for NorA. Values are plotted as a function of log₂[EtBr].

**Figure 4.**
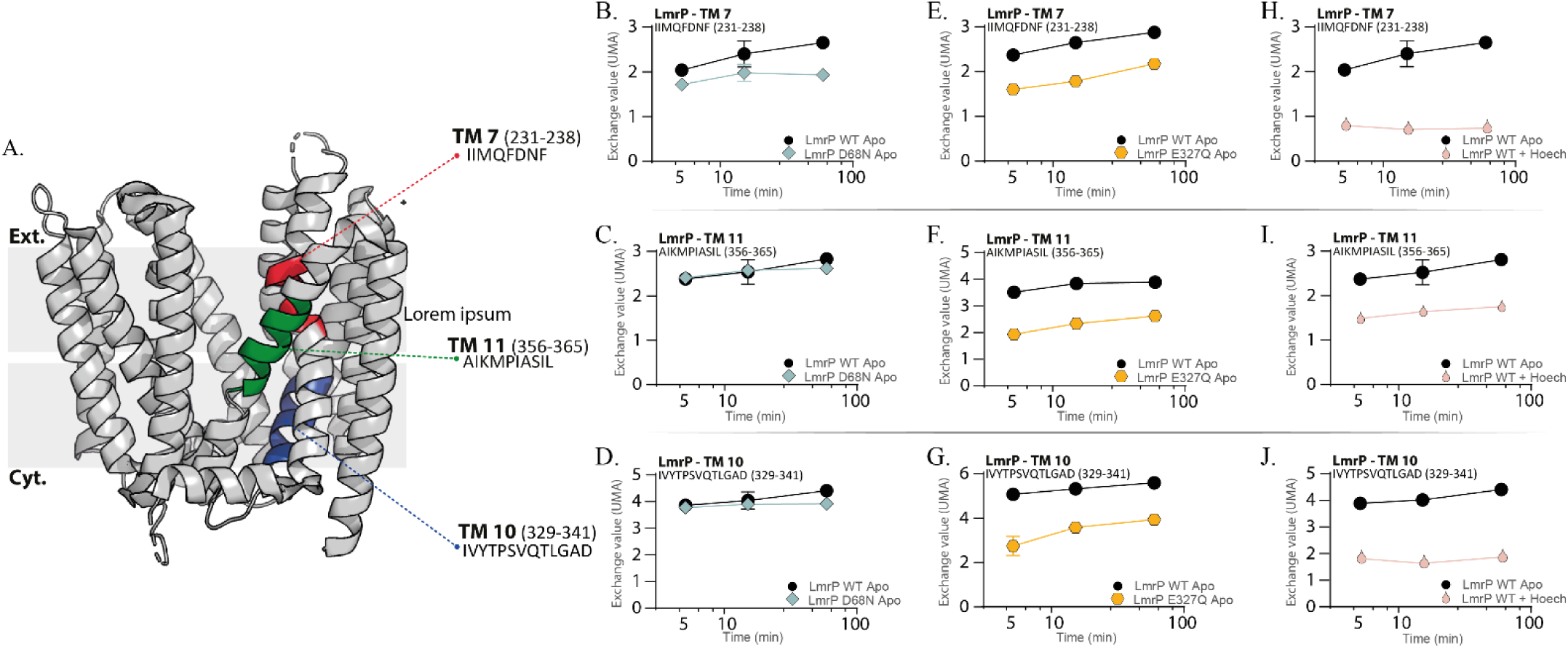
The E327Q mutation recapitulates the effect of substrate binding whereas D68N does not. **(A)** Location of the three analysed peptides mapped onto the LmrP crystal structure (PDB: 6T1Z): TM7 peptide 231–238, TM10 peptide 329–341 and TM11 peptide 356–365. **(B-J)** Representative deuterium uptake plots for the indicated peptides, comparing LmrP WT apo with LmrP D68N apo (**B-D**), LmrP E327Q apo (**E-G**), or LmrP WT in the presence of Hoechst 33342 (**H-J**). LmrP WT apo is shown as black circles, LmrP D68N apo as light-green diamonds, LmrP E327Q apo as orange hexagons and LmrP WT + Hoechst33342 as salmon teardrops. Data are shown as mean ± SD across three replicates (*n* = 3).

Previous HDX-MS studies of non-multidrug MFS transporters, including the bacterial sugar transporters XylE, MelB and LacY, did not detect comparable exchange within transmembrane helices ^32,33,36,37^. In those systems, uptake was restricted to peptides that partially or fully encompassed loop regions, while peptides entirely contained within α-helices showed little, if any, exchange, remaining below 10% maximum R.F.U. even at the longest labeling times (**Supplementary Fig. 5**). This clear difference suggests that transmembrane helix flexibility is not a general feature of MFS transporters, but may instead be associated with the polyspecific ligand recognition that defines multidrug efflux pumps.

### The same substrate differentially modulates the dynamics of flexible helices

To understand how substrate binding affects the local structural dynamics of MFS MDR transporters, we performed measurements in the presence ethidium bromide (EtBr), a common substrate for the three different transporters^27,31,38–40^. We expressed the differences in deuterium uptake as changes in the relative fractional uptake (ΔR.F.U.), which is positive when regions take up more deuterium in presence of EtBr (relative to the apo) or negative when they are protected from uptake. For the three transporters, analysis of the ΔR.F.U heatmaps shows different structural responses across the protein sequences, with the most important changes occurring in the helices identified as flexible (**Fig. 2A, E, G**). Notably, both the extent and the direction of the ΔR.F.U changes (increased or decreased exchange) vary substantially across the different proteins.

Ethidium binding to LmrP leads to a protection from exchange that localizes on the C-lobe, indicating a reduction in local structural dynamics (**Fig. 2A**). A breakdown into individual peptides of the aggregated heatmap is presented for the 15-minutes time-point, allowing to observe that transmembrane peptides identified as intrinsically dynamic on TMH7, TMH10, and TMH11 display significant protection from exchange, with ΔR.F.U. values of individual peptides up to 25% (**Fig. 2A-E and Supplementary Fig. 6**). This indicates protection from exchange and a decrease in flexibility in these specific regions. A model of ethidium bound LmrP predicts direct interaction between these three TMHs and the substrate (**Fig. 2E** and **Supplementary Fig. 9A**). In the case of NorA, an overall protective effect is observed on the ΔHDX heatmaps upon ethidium binding (**Fig. 2F**). The effect is much less pronounced compared to LmrP, with the highest ΔR.F.U. values not exceeding 10% (**Fig. 2F-H)**. TMH4 and TMH5, identified as flexible, display significant protection upon ethidium binding. (**Fig 2F-H and Supplementary Fig. 7**)^42,43^. This protection indicates that the substrate effectively stabilizes the surrounding helices, reducing their local structural dynamics. This data is consistent with a model of NorA bound to ethidium (**Fig. 2H** and **Supplementary Fig.9B**), that predicts direct interactions with TMH5, especially with residues N137 and F140, but not with TMH4, suggesting that the observed changes in local structural dynamics are a combination of direct and allosteric effects. In contrast, ΔHDX data of QacA reveals a widespread increase in deuterium exchange across nearly the entire protein scaffold upon ethidium binding (**Fig. 2I-K** and **Supplementary Fig. 8**). indicating increase in overall structural dynamics, with the strongest effect in the helices identified as flexible. This deprotection is particularly pronounced in TMHs 2, 5 and 10, and is consistent over biological replicates. Recently reported experimental structures of wild-type QacA in the apo state and bound to ethidium, obtained by X-ray crystallography, reveal direct interactions with specific residues in TMHs2, 5, and 13 ^31^ (**Supplementary Fig. 9C**). Comparison of these two states further highlights a pronounced ligand-induced kink in TMH5, indicative of a local change in secondary structure that is consistent with our experimental observations (**Supplementary Fig. 10**). By contrast, no direct interaction is observed with TMH10, suggesting that the increased exchange detected in this helix may arise from an allosteric effect. Positioned at the interface between the N- and C-lobes, TMH10 is well placed to participate in conformational rearrangements.

Together, these findings identify intrinsically flexible helices as key sites of ligand-dependent modulation of local structural dynamics, while also showing that the direction and magnitude of this modulation is transporter-dependent. Notably, the observed changes in structural dynamics extend beyond the binding site, consistent with a broader allosteric response of the transporter to ligand binding.

### Changes in local dynamics are accompanied by changes in global thermostability

To determine whether the ligand-dependent changes in local structural dynamics are associated with changes in the overall thermal stability of the transporters, we next performed nano differential scanning fluorimetry (nanoDSF) measurements. Whereas HDX-MS provides a localized view of the effect of ethidium binding to local structural dynamics, nanoDSF offers an orthogonal readout of its net effect on global protein stability.

Because nanoDSF monitors changes in intrinsic tryptophan fluorescence during thermal unfolding, interpretation of the resulting profiles depends on both the number and structural distribution of tryptophan residues in each transporter. The three transporters differ in both the number and distribution of tryptophan residues, with NorA containing three, LmrP seven, and QacA nine tryptophans located in both solvent-exposed and solvent-shielded environments (**Fig. 3A, B and C**). Consistent with this, their thermal unfolding profiles differed markedly (**Fig. 3D, E and F**).

In particular, LmrP displayed a multiphasic profile with an intermediate plateau, precluding reliable determination of melting temperatures using standard derivative-based or sigmoidal fitting methods^44–46^. In contrast, NorA and QacA yielded unfolding curves compatible with conventional nanoDSF analysis. However, to ensure consistency across the three transporters, we quantified ligand effects using the apparent transition temperature (T_app_) determined by geometric analysis of the fluorescence ratio curves (**Methods**)^47^.For LmrP, analysis of the curve showed a clear transition at 50.2 °C in the apo condition that shifted to 55.4 °C in the presence of a saturating amount of EtBr (**Fig. 3D**), indicating thermostabilization upon binding. The observed stabilization was validated across multiple biological replicates, yielding a mean ΔT_app_ of 5.2 ± 2.3°C (**Supplementary Fig. 11**). EtBr binding shifts the melting temperature of NorA from 51.7 °C to 56.6 °C, a gain of nearly 4.2 °C ± 1.0°C, again indicating thermostabilization upon binding (**Fig. 3E and Supplementary Fig.12**). For QacA, two transition events were clearly visible in the unfolding curve (**Fig. 3F**). The first apparent transition temperature (T_app1_) was around 47.5 °C in both the apo and ethidium-bound conditions. The second apparent transition temperature (T_app2_) decreased from 70.3 °C in the apo condition to 67.2 °C in the bound condition, indicating thermal destabilization, with a ΔT_app2_ of 3.1 ± 1.0°C (**Supplementary Fig. 13**).

This ligand-induced destabilization is further confirmed by a titration experiment showing that T_app2_ progressively decreases with increasing EtBr concentrations (**Fig. 3G**). A similar titration experiment was performed for NorA and confirmed the EtBr-induced thermostabilization, as shown by the gradual increase in T_app_ over increasing ethidium concentrations (**Fig. 3H**). Overall, the nanoDSF measurements show that binding of the same substrate stabilizes LmrP and NorA, but destabilizes QacA.

Together, the HDX-MS and nanoDSF data show that a shared substrate can remodel both local structural dynamics and global thermal stability in different ways across related MFS multidrug transporters. Thus, even within a conserved fold energized by the same proton motive force, substrate recognition is coupled to distinct energetic and dynamical responses.

### Protonation-substrate coupling controls helix dynamics in LmrP

Having examined the effect of substrate binding, we next asked how the other driving force in the transport cycle, that is the proton motive force, influences local structural dynamics. Specifically, in proton-coupled MFS multidrug transporters, sequential protonation of conserved acidic residues has been shown to drive the structural transitions required for substrate extrusion ^23,27,38^. LmrP provides a useful system for this analysis because its conformational landscape, and its modulation by pH and protonation mimicking mutants, has been well defined by DEER^23,48^ and smFRET^49^ spectroscopies.

Apo wild-type LmrP at pH 7 predominantly adopts an outward-open conformation^23,49^. Residue D68 is part of the most conserved motif in the MFS family – coined motif A – and is part of charge relay network that stabilizes the outward-open conformation^5,50,51^. The D68N mutation is a useful mimic of protonation and shifts the equilibrium toward the inward-facing state^23,48^. E327Q mutant is informative for a different reason: Conserved residue E327 is involved both in proton coupling as the staring point of the proton translocation pathway ^52–54^ and in direct ligand interactions^24^, including with ethidium^52^. The E327Q mutation preserves the outward-facing ensemble under conditions that otherwise promote inward-facing state stabilization^23^. Together, these state- and site-specific perturbations provide a framework to interpret HDX-MS changes in relation to both global conformational rearrangements and local mechanistic effects. We therefore performed HDX-MS measurements on the protonation-mimicking D68N and E327Q mutants, focusing our analysis on the flexible transmembrane helices identified above, TMHs 7, 10 and 11, to determine whether their local conformational plasticity is modulated by proton binding and how this relates to the global conformational landscape of LmrP (**Fig. 4**).

We found that the intrinsic flexibility of TMHs 10 and 11 was preserved in the D68N mutant (**Fig. 4C, D**), despite its global conformational shift toward the inward-facing (IF) state, as confirmed by the characteristic ΔHDX pattern observed in the N-lobe (**Supplementary Fig. 14**). These data indicate that the structural flexibility of these helices is a constitutive feature of LmrP and is largely independent of whether the transporter populates outward- or inward-facing conformations.

In contrast, ΔHDX-MS analysis of the E327Q mutant revealed marked protection from exchange in TMHs 7, 10 and 11, with an approximately 25% reduction in R.F.U. compared with the wild type (**Fig. 4E-G** and **Supplementary Fig. 15**). Strikingly, this pattern mirrors the protection observed upon ethidium binding (**Fig. 2A-E**). The fact that a conservative E to Q substitution mirrors the effect of ethidium binding is particularly informative: it shows that the HDX protection signature does not require the physical presence of the substrate molecule itself, but can be triggered by neutralizing a single conserved acidic residue. The ability of this single side-chain substitution to recapitulate the HDX signature induced by a bulky substrate argues against a simple solvent-shielding effect caused by substrate occupancy.

This observation led us to hypothesize that the loss of flexibility in these TMHs is directly controlled by the charge state or binding status of E327. In other words, interaction of E327 with either a proton or a ligand may rigidify the flexible helices TMHs 7, 10 and 11. To test this hypothesis, we performed additional ΔHDX-MS measurements in the presence of Hoechst 33342, as the crystal structure of LmrP bound to Hoechst 33342 shows a direct interaction between the ligand and E327. Consistent with our hypothesis, all three helices displayed protection from exchange in the presence of Hoechst 33342 (**Fig. 4H-J and Supplementary Fig. 16**). To further assess the generality of this mechanism, we tested another substrate, the antibiotic tetracycline, and again observed protection from exchange in the flexible helices (**Supplementary Fig. 16**).

Together, these results identify E327 as a coupling site that links protonation and substrate binding to local helix rigidification within the C-lobe. This suggests that energy coupling in LmrP is mediated, at least in part, through local modulation of helix flexibility within the broader alternating-access framework.

### A revised transport model links protonation and ligand binding to local plasticity in LmrP

To illustrate our findings, we can now propose an updated transport cycle for the prototypical MFS MDR LmrP that explicitly integrates the role of local helix dynamics (**Fig. 5**), and in particular how protonation or ligand binding at the conserved E327 glutamate acts as a switch for helical flexibility. In its resting apo state, TMHs 7, 10, and 11 in the C-lobe are flexible and do not always adopt a fully helical conformation, thereby priming the transporter for ligand binding (**Fig. 5A**). The substrate comes in and, through its cationic moiety, interacts with E327 on TMH10, inducing local rigidification of this region and activating an allosteric network that also rigidifies the neighboring helices TMHs 7 and 11 (**Fig. 5B**). This is accompanied by a conformational transition toward an outward-open state, as previously shown by DEER spectroscopy with several substrates^24^ (**Fig. 5C**). Competition between extracellular protons and substrate for E327 promotes substrate release to the extracellular medium (**Fig. 5D**). Subsequent proton translocation from E327 to other acidic residues restores flexibility within the transmembrane helices. Protonation of D68 then triggers a conformational switch that primes the transporter for a new transport cycle^23,48^, with the extracellular side closed and local helical flexibility restored (**Fig. 5E**).

**Figure 5.**
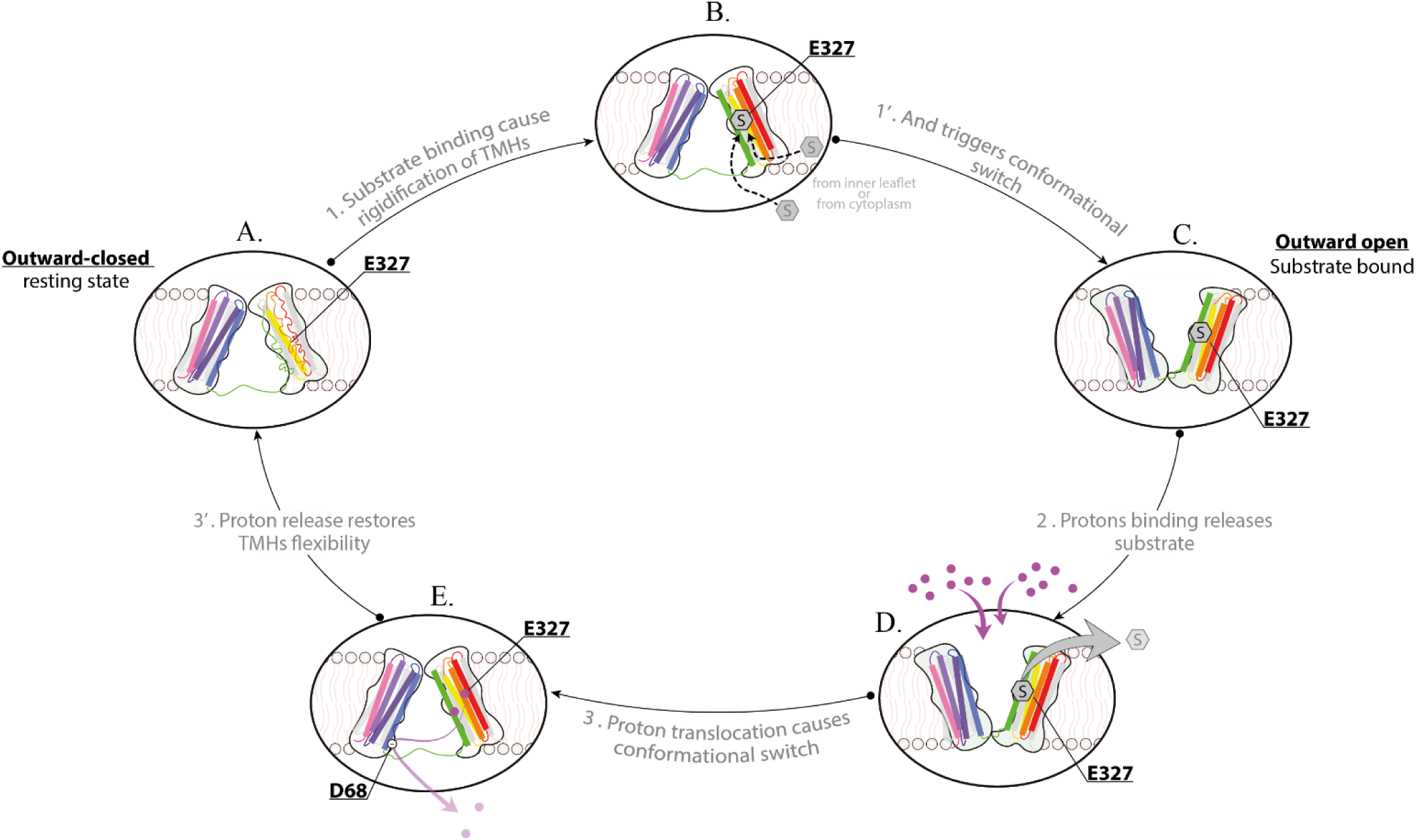
Proposed transport mechanism of LmrP driven by local helix dynamics. (A) in the resting state, C-lobe TMs 7, 10 and 11 exhibit high flexibility. (B) The cationic substrate binding to E327 induces local rigidification of the helices. (C) The structural consolidation facilitates the transition to an outward-open conformation, consistent with the DEER data. (D) Under the proton motive force, extracellular protons compete with the substrate, triggering the release. (E) The proton translocation from E327 to other acidic residues, including D68, triggers the conformational reset.

## Discussion

The systematic comparison of three homologous multidrug efflux pumps reveals that specific transmembrane helices are intrinsically flexible and that this flexibility is modulated by ligand binding, whether by protons or substrates. More broadly, this work shows that dynamic fingerprinting can uncover an additional layer of regulation in membrane proteins, beyond what is apparent from static structures alone. This is particularly well illustrated by LmrP, where analysis of protonation-mimic mutants and multiple ligands shows that local helix flexibility provides a mechanistic link between substrate recognition, proton coupling and conformational switching.

NMR played a central role in establishing that local dynamics are key determinants of protein function^55–57^. Elegant studies on soluble proteins demonstrated in exquisite detail how changes in local dynamics and secondary structure are directly linked to function, for example in enzymatic catalysis^58,59^. By contrast, the technical challenges associated with applying NMR to membrane proteins have limited comparable insights for this important class of proteins^60–62^. As a result, our molecular understanding of membrane protein function has emerged largely from high-resolution structures, supported by biochemical and biophysical approaches^62^. HDX–MS does not (yet) offer the same level of resolution as NMR, but it is far more accessible for these challenging targets. In particular, improvements in sensitivity, automation, and data-analysis workflows have made it possible to use HDX–MS to dissect the molecular mechanisms of many transporters, initially as a complementary biophysical tool, ^63,64^, but increasingly as an approach capable of providing unique mechanistic insight1^16,65–67^ Integrating local dynamics into molecular mechanisms is a necessary next step for transporters, building directly on existing structural models.

Recent studies have serendipitously revealed local flexibility in transporters and highlighted its functional consequences. For example, a study on the archaeal glutamate transporter Glt_Ph_ reported heterogeneous substrate binding, manifested as a biphasic affinity curve^68^. Detailed analysis of cryo-EM data from a Glt_Ph_ mutant restricted to the outward-open state showed that this state comprises multiple subconformations, which differ primarily in the degree of “unpacking” of specific transmembrane helices. Importantly, the heterogeneous transport kinetics were shown to arise from these local subconformations coexisting within a single global conformational state. Extensive work on NSS family transporters, particularly on the archaeal NSS homolog LeuT, has established helical unwinding as a necessary step in transitions between conformational states^16,67,69,70^. This unwinding is itself regulated by ion and substrate binding, with the concomitant presence of leucine and Na⁺ suppressing unwinding compared with apo or K⁺-bound conditions. Together, these findings establish local helical flexibility as an important component of transporter molecular mechanisms. The next challenge is to design experiments that directly connect this flexibility to transport function^71^.

Another finding from the present work is that ligand-induced modulation of local structural dynamics is protein-specific. The similarity of the protein folds studied here as well as their identical proton-coupled antiport mechanism made a broadly similar structural response in the presence of an identical substrate a plausible scenario. Instead, each transporter presents a distinct pattern of hydrogen-bond network rearrangements upon binding. In LmrP, substrate binding and neutralization of E327 converge on protection of TMHs 7, 10 and 11, consistent with local rigidification of a flexible C-lobe region. In NorA, ligand-induced effects are more restricted, whereas QacA shows a broader increase in exchange, indicative of enhanced flexibility rather than rigidification. These results show that a conserved fold can support multiple sequence-encoded mechanistic solutions. In other words, while the fold defines the structural framework and is a good predictor of the global function (e.g. “this is a transporter”), the sequence determines how that framework is dynamically and energetically exploited at the molecular level (e.g. “this is a multidrug efflux pump, that relies on a conserved glutamate on TMH10 to couple proton translocation to substrate extrusion”).

More broadly, this work shifts the focus from static structural descriptions toward a dynamic, sequence-encoded view of transport mechanisms. This dynamic view provides a mechanistic framework for interpreting deep mutational scanning studies^10,11^, in which residues distant from the ligand-binding site can nevertheless reshape polyspecific transport through allosteric effects on the conformational ensemble. How these dynamics are tuned by the native membrane environment, including lipid composition, electrochemical gradients, molecular crowding, and membrane curvature, will allow to correlate the complexity of cellular function to molecular mechanism. Our results also underscore the need for structural models that explicitly incorporate dynamics, an emerging goal of next-generation computational and generative approaches, still very much in its infancy^71–73^. Integrating experimentally validated dynamic constraints into such models will be essential for predictive mechanistic understanding and for the rational design of selective modulators targeting conformational landscapes.

## Methods

Detergents were purchased from Anatrace. Unless otherwise stated, all other chemicals were purchased from Sigma-Aldrich, now Merck.

### LmrP Expression

As previously described^24^, the *lmrp* gene was cloned in the pHLP5-3C expression plasmid in a construction containing the *lmrp* gene, 3C cleavable sequence and a C terminal 6-his tag. The D68N mutant and E327Q mutant constructions were generated by single point mutagenesis using the QuikChange Lightning cloning kit (Stratagene). Plasmids were transformed into *L. lactis* strain NZ9000 through electroporation (adapted from ^74^), and a single colony was used for protein expression. Cells were grown at 30°C in M17 medium (Biokar) complemented with 0.5% (wt/vol) glucose and 5 µg.mL^−1^ chloramphenicol until the optical density at 600 nm (OD_600_) reached 0.8. Overexpression of LmrP was then induced by the addition of 1:1,000 of the supernatant of the nisin-producing *L. lactis* strain NZ9700. After 2h of further incubation still at 30°C, cells were collected by centrifugation at 4,000g.

### QacA Expression

*The E. coli* BW25113 *Δ* AcrB strain, previously transformed with a pBAD30-QacA-6His was used to overexpress the protein QacA. Cells were grown in 20 mL preculture overnight at 37°C, at 160 RPM in LB medium supplemented with 100 µg.mL^−1^ ampicillin. Then, preculture is mixed with 980 mL of TB medium also supplemented with 100 µg.mL^−1^ ampicillin. The overexpression was induced using 0.02% (w/v) L-arabinose when OD_600_ reached 0.7. After 4h of further incubation, now at 30°C, cells were collected by centrifugation at 4,000g.

### NorA Expression

*Escherichia coli* BL21 strain, have been previously transformed with a pTrcHis2C-NorA-His6. Plasmid was provid by Davud Hoper. to overexpress the protein NorA. Cells were grown in 20 mL preculture overnight at 37°C, at 160 RPM in TB medium supplemented with 100 µg.mL^−1^ ampicillin. Then, preculture is mixed with 980 mL of TB medium at 37°C, at 160 RPM supplemented with 100 µg/mL ampicillin. The expression was induced at OD_600_ = 0.7 using 0.5 mM of IPTG. After 4h of further incubation, now at 30°C, cells were collected by centrifugation at 4,000g.

### Proteins purification and characterization

Cell pellets were washed once and resuspended in buffer prior to lysis. L. lactis cells were resuspended in 50 mM HEPES, pH 7.0, while E. coli cells were resuspended in 20 mM HEPES, 300 mM NaCl, pH 7.0. Cell suspensions were flash-frozen and stored at −80 °C until further use.

L. lactis cells were incubated with lysozyme (10 mg.mL⁻¹ of resuspended cells) for 1 h at 30 °C and DNase I (20 U.mL⁻¹ of culture) with his cofactor MgCl_2_ (10 mM final) for 15 min also at 30 °C, followed by disruption using a French press at 30,000 psi. E. coli cells were incubated with DNase I under identical conditions and disrupted using a French press at 17,500 psi. Cell debris was removed by centrifugation (2 × 20 min at 10,000 g).

Membrane fractions were isolated by ultracentrifugation at 100,000 g for 2 h at 4 °C. Membrane pellets were resuspended in solubilization buffer consisting of either 50 mM HEPES, 150 mM NaCl, 10% (v/v) glycerol, pH 7.0 (LmrP), or 50 mM HEPES, 300 mM NaCl, 10% (v/v) glycerol, pH 7.0 (QacA and NorA). Membranes were solubilized by incubation with 1.2% (w/v) *β*-DDM for 2 h at 4°C under gentle agitation. Insoluble material was removed by ultracentrifugation at 100,000 g for 30 min.

Solubilized proteins were purified by immobilized metal affinity chromatography using Qiagen Ni-NTA agarose resin. After binding for 2 h at 4 °C, the resin was washed with solubilization buffer containing 0.05% *β*-DDM and either 30 mM (LmrP) or 50 mM (QacA and NorA) imidazole. Proteins were eluted with solubilization buffer containing 300 mM (LmrP) or 350 mM (QacA and NorA) imidazole.

Eluted proteins were desalted using PD-10 columns (GE Healthcare) and further purified by size-exclusion chromatography using a Superdex 200 Increase 10/300 column (**Supplementary Fig. 17**). The final SEC buffer consisted of 25 mM (LmrP) or 20 mM (QacA and NorA) HEPES, 100 mM (LmrP) or 150 mM (QacA and NorA) NaCl, 0.02% *β*-DDM, pH 7.0.

### Hydrogen Deuterium eXchange - Mass Spectrometry

Hydrogen-deuterium exchange mass spectrometry (HDX-MS) experiments were performed on Waters Synapt G2 or Synapt G2-Si electrospray ionization quadrupole time-of-flight ion mobility mass spectrometers. Measurements were conducted at the Martens Laboratory (ULB), MSLab (ULiège), and the Biomolecular Mass Spectrometry Facility (University of Leeds).

Membrane proteins samples were prepared at a concentration of 40 to 60 µM using Vivaspin concentrators with 50 kDa MWCO. To standardize sample conditioning, all samples were pre-incubated on ice for 30 min, either in the absence or presence of substrate. When applicable, substrates were added at a 1:30 protein-to-substrate molar ratio.

HDX labeling was initiated by diluting 7 µL of protein sample into 63 µL of labeling buffer (same composition as the SEC buffer described in the *Protein purification and characterization* section, except prepared in D₂O). Labeling reactions were performed at 20 °C using a plate heater and allowed to proceed for defined time intervals ranging from 30 s to 1 h. Labeling reactions were quenched by the manual addition of an equal volume of pre-chilled quench buffer (50 mM Phosphate buffer, 1.0% formic acid, 0.05% *β*-DDM, pH 2.3).

Following quenching, samples were flash-frozen in liquid nitrogen and stored at −80 °C until HDX-MS analysis. Prior to injection, samples were thawed manually by holding them at room temperature, with a controlled and reproducible thawing time of approximately 75 s.

Once thawed, the samples were digested online on Enzymate BEH Pepsin Column (Waters Corporation) at 200 µl.min^−1^ and at 20 °C with a pressure 3 kPSI (10 kPSI when setups were equipped with a Back Pressure Regulator). Peptic peptides were trapped for 3 min on an Acquity UPLC BEH C18 VanGuard Pre-column (Waters Corporation) at a 200 µl.min^−1^ flow rate in water (0.1% formic acid in HPLC water pH 2.5) before eluted to an Acquity UPLC BEH C18 Column for chromatographic separation. Separation was done with a linear gradient buffer (3–45% gradient of 0.1% formic acid in acetonitrile) at a flow rate of 40 µl.min^−1^. Peptides identification and deuteration uptake analysis was performed on the Synapt G2 Si in ESI ± MSE mode (Waters Corporation). Leucine enkephalin was applied for mass accuracy correction and sodium formate was used as calibration for the mass spectrometer. MS^E^ data were collected by a 20-45 eV transfer collision energy ramp. The pepsin column was washed between injections using pepsin wash buffer (1.5 M Guanidinium HCl, 4% (v/v) acetonitrile, 0.8% (v/v) formic acid). A cleaning run was performed for each three sample to prevent peptide carry-over. All deuterium time points were performed in technical triplicate.

### Data evaluation and statistical analysis

Acquired reference MS^E^ data were analyzed by PLGS (ProteinLynx Global Server 3.0.2, Waters) to identify the peptic peptides, then all the MS^E^ data including reference and deuterated samples were filtered and processed by DynamX v.3.0 (Waters) for deuterium uptake determination with the following filtering parameters: minimum intensity of 1000, minimum and maximum peptide sequence length of 5 and 25, respectively, minimum MS/MS products of 2, minimum products per amino acid up to 0.30, minimum score of 5, and a maximum MH^+^ error threshold of 15 p.p.m.. Peptide identification was considered valid only when a peptide was consistently observed in a minimum of three out of five independent reference samples.

When further statistical analysis was required, Deuteros 2.0 (v 2.4.2), developed by Lau et al.^43^, was used. Differences between two states is first evaluated using the hybrid significance test, which assesses whether observed deuterium uptake differences exceed a threshold defined by the selected confidence level (98% confidence; *α* = 0.02). Statistical significance is subsequently confirmed using Welch’s t-test.

### Dataset division based on 3D structures

To perform and fulfill the analysis of our data on LrmP, NorA and QacA, we divided the dataset based on secondary structure elements, distinguishing between transmembrane helices and loop regions. Helices were defined as segments consistently structured in both conformations of three-dimensional models of LmrP: the outward-open conformation (PDB ID: 6T1Z) and the inward-open model predicted by AlphaFold^75^, of NorA: the outward-open conformation (PDB ID: 7LO7 and 7LO8) and the inward-open conformation (PDB ID: 8TTE) and of QacA: the outward-open conformation (PDB ID: 7Y58) and the inward-open conformation model predicted by AlphaFold. Peptides were assigned to the helix dataset only if all residues were fully contained within well-defined *α*-helical regions; any peptide with one or more residues located outside these regions was assigned to the loop dataset.

### HDX-MS data analysis on subdivided datasets (helices vs loops)

The structural segmentation yielded two distinct datasets: the helix dataset, comprising peptides from canonical transmembrane helices, and the loop dataset, encompassing both intrinsically disordered regions and helix-adjacent segments lacking stable secondary structure. The rationale behind this division is grounded in HDX theory: loop regions, being solvent-exposed and not stabilized by backbone hydrogen bonds, are expected to exhibit more rapid deuterium exchange than structured helical regions, which are typically buried and hydrogen-bonded.

To derive a quantitative criterion for classifying peptides as flexible or rigid, we used the mean R.F.U. value of the loop dataset as a reference threshold. Peptides exhibiting mean R.F.U. values above this threshold were classified as flexible.

### NanoDSF Thermostability measurements

Thermal denaturation was monitored by recording intrinsic tryptophan fluorescence at 330 nm and 350 nm, which reflects changes in the local environment of tryptophan residues during unfolding. Monitoring fluorescence at both wavelengths allows detection of emission peak shifts while reducing background contributions. Purified LmrP WT and mutants, QacA and NorA were diluted to a final concentration of 5 to 10 µM in their respective SEC buffer, containing HEPES, NaCl, Glycerol and *β*-DDM. For apo conditions alone, the proteins were thawed on ice for 15 min. For substrate bound conditions, the substrates were prepared initially prepared at 100 mM in DMSO then diluted to 25 mM in SEC buffer (not required for EtBr which is water soluble) and finally diluted again to final concentration of 1 mM. Then substrates were incubated in a ratio of 1:25 (5 µM protein : 125 µM ligand) on ice for 30 min. 8–10 µL of the prepared protein samples were transferred into standard grade Prometheus NT.48 capillaries. The PR.ThermControl (version 2.1.2) software was used to set up a melting scan from 15°C to 90°C with a ramp rate of 1°C/min. The fluorescence emission at 330 nm (F330), 350 nm (F350), the 350:330 ratio, and the light scattering signal were recorded over the course of the melting scan. The effect of all ligands on protein stability was tested in the same run for each protein and compared to the reference apo sample without lipid added (apo). Initial fluorescence was checked as some substrates are fluorescent in the range of tryptophan emissions. Only substrates maintaining the initial fluorescence above the threshold of 1000 mAU were conserved. We systematically performed technical duplicate.

### NanoDSF data analysis

Because several nanoDSF unfolding profiles were non-sigmoidal and/or lacked well-defined pre-and post-transition plateaus, apparent transition temperatures were extracted using a geometrical tangent-extrapolation procedure. The apparent transition temperature (T_app_) was estimated from sigmoidal thermal unfolding curves using the tangent method. This graphical approach was selected for its robustness against experimental noise, offering an advantage over first-derivative analysis which tends to amplify fluctuations in point-to-point slope changes. Briefly, parallel tangents were constructed at the onset of the pre-transition and post-transition regions, corresponding to the points where the slope of the unfolding curve initially deviates from linearity. A median line was subsequently drawn equidistant to and parallel with these two tangents. The intersection point of this median line with the experimental unfolding curve was identified as the inflection point, providing the estimate for T_app_. T_app_ values were used as operational descriptors of the unfolding profiles rather than as thermodynamic melting temperature. Data analysis was performed using GraphPad Prism.

## Acknowledgments

We thank Jean-Marie Ruysschaert, George Nyasha Chiduza and the members of the Govaerts Lab and of the Martens lab for discussions and feedback throughout this work; We thank the Mass spectrometry Facility from University of Leeds, especially the facility manager James Ault. We also thank the Mass Spectrometry Laboratory at University of Liège for access to instrumentation and Gabriel Mazzuchelli and Johann Far for expert technical support and assistance with data acquisition. The authors acknowledge the use of EMBL Hamburg NanoTemper Prometheus NT.48 though MOSBRI proposal and thank the supporting team for their help, specifically Angelica Struve Garcia and Stephan Niebling. We thank Prof. Melissa Brown for providing the plasmids for QacA expression and David Hooper for providing the plasmid for NorA expression.

## Authors contribution

K.C.W. and C.M. conceived the study and designed the experiments. K.C.W. and M.M performed the expression, purification of proteins prior HDX-MS experiments. K.C.W. carried out the HDX-MS experiments with the help of R.M. and T.T. K.C.W. analyzed the HDX-MS data. K.C.W. carried out the nanoDSF experiments and performed the data analysis. K.C.W. and C.M. wrote the paper with input from all the authors.

## References

1. Reygaert, W. C. An overview of the antimicrobial resistance mechanisms of bacteria. AIMS Microbiol. 4, 482–501 (2018).

2. Piddock, L. J. V. Multidrug-resistance efflux pumpsnot just for resistance. Nat. Rev. Microbiol. 4, 629–636 (2006).

3. Du, D. et al. Multidrug efflux pumps: structure, function and regulation. Nat. Rev. Microbiol. 16, 523–539 (2018).

4. Blair, J. M. A., Webber, M. A., Baylay, A. J., Ogbolu, D. O. & Piddock, L. J. V. Molecular mechanisms of antibiotic resistance. Nat. Rev. Microbiol. 13, 42–51 (2015).

5. Drew, D., North, R. A., Nagarathinam, K. & Tanabe, M. Structures and General Transport Mechanisms by the Major Facilitator Superfamily (MFS). Chem. Rev. 121, 5289–5335 (2021).

6. Kumar, S. et al. Functional and Structural Roles of the Major Facilitator Superfamily Bacterial Multidrug Efflux Pumps. Microorganisms 8, 266 (2020).

7. Schindler, B. D. & Kaatz, G. W. Multidrug efflux pumps of Gram-positive bacteria. Drug Resist. Updat. Rev. Comment. Antimicrob. Anticancer Chemother. 27, 1–13 (2016).

8. Fluman, N. & Bibi, E. Bacterial multidrug transport through the lens of the major facilitator superfamily. Biochim. Biophys. Acta BBA - Proteins Proteomics 1794, 738–747 (2009).

9. Yee, S. W. et al. The full spectrum of SLC22 OCT1 mutations illuminates the bridge between drug transporter biophysics and pharmacogenomics. Mol. Cell 84, 1932–1947.e10 (2024).

10. Meier, G. et al. Deep mutational scan of a drug efflux pump reveals its structure–function landscape. Nat. Chem. Biol. 1–11 (2022) doi:10.1038/s41589-022-01205-1.

11. Miller, S. T., Henzler-Wildman, K. A. & Raman, S. Energetic and structural control of polyspecificity in a multidrug transporter. Proc. Natl. Acad. Sci. U. S. A. 122, e2511892122 (2025).

12. Kuzmanic, A., Pannu, N. S. & Zagrovic, B. X-ray refinement significantly underestimates the level of microscopic heterogeneity in biomolecular crystals. Nat. Commun. 5, 3220 (2014).

13. Tang, W. S., Zhong, E. D., Hanson, S. M., Thiede, E. H. & Cossio, P. Conformational heterogeneity and probability distributions from single-particle cryo-electron microscopy. Curr. Opin. Struct. Biol. 81, 102626 (2023).

14. Wang, L. C., Krishnamurthy, S. & Anand, G. S. Hydrogen Exchange Mass Spectrometry Experimental Design. in Hydrogen Exchange Mass Spectrometry of Proteins 19–35 (2016). doi:10.1002/9781118703748.ch2.

15. Vinciauskaite, V. & Masson, G. R. Fundamentals of HDX-MS. Essays Biochem. 67, 301–314 (2023).

16. Merkle, P. S. et al. Substrate-modulated unwinding of transmembrane helices in the NSS transporter LeuT. Sci. Adv. 4, eaar6179 (2018).

17. Wright, P. E. & Dyson, H. J. Intrinsically unstructured proteins: re-assessing the protein structure-function paradigm. J. Mol. Biol. 293, 321–331 (1999).

18. Katta, V., Chait, B. T. & Carr, S. Conformational changes in proteins probed by hydrogen-exchange electrospray-ionization mass spectrometry. Rapid Commun. Mass Spectrom. 5, 214–217 (1991).

19. Engen, J. R. Analysis of Protein Conformation and Dynamics by Hydrogen/Deuterium Exchange MS. Anal. Chem. 81, 7870–7875 (2009).

20. Yu, J.-L., Grinius, L. & Hooper, D. C. NorA Functions as a Multidrug Efflux Protein in both Cytoplasmic Membrane Vesicles and Reconstituted Proteoliposomes. J. Bacteriol. 184, 1370–1377 (2002).

21. Mitchell, B. A., Brown, M. H. & Skurray, R. A. QacA Multidrug Efflux Pump from Staphylococcus aureus: Comparative Analysis of Resistance to Diamidines, Biguanidines, and Guanylhydrazones. Antimicrob. Agents Chemother. 42, 475–477 (1998).

22. World Health Organization. WHO Bacterial Priority Pathogens List, 2024: Bacterial Pathogens of Public Health Importance to Guide Research, Development and Strategies to Prevent and Control Antimicrobial Resistance. (Geneva, 2024).

23. Masureel, M. et al. Protonation drives the conformational switch in the multidrug transporter LmrP. Nat. Chem. Biol. 10, 149–155 (2014).

24. Debruycker, V. et al. An embedded lipid in the multidrug transporter LmrP suggests a mechanism for polyspecificity. Nat. Struct. Mol. Biol. 27, 829–835 (2020).

25. Maklad, H. et al. Antibiotics-induced conformational heterogeneity of a multidrug transporter revealed by single-molecule FRET. 2025.05.17.654648 Preprint at 10.1101/2025.05.17.654648 (2025).

26. Brawley, D. N. et al. Structural basis for inhibition of the drug efflux pump NorA from Staphylococcus aureus. Nat. Chem. Biol. 18, 706–712 (2022).

27. Li, J. et al. Proton-coupled transport mechanism of the efflux pump NorA. Nat. Commun. 15, 4494 (2024).

28. Quistgaard, E. M., Löw, C., Guettou, F. & Nordlund, P. Understanding transport by the major facilitator superfamily (MFS): structures pave the way. Nat. Rev. Mol. Cell Biol. 17, 123–132 (2016).

29. Reddy, V. S., Shlykov, M. A., Castillo, R., Sun, E. I. & Saier Jr, M. H. The major facilitator superfamily (MFS) revisited. FEBS J. 279, 2022–2035 (2012).

30. Majumder, P. et al. Cryo-EM structure of antibacterial efflux transporter QacA from Staphylococcus aureus reveals a novel extracellular loop with allosteric role. EMBO J. 42, e113418 (2023).

31. Jodaitis, L. et al. Structural basis of drug efflux by the staphylococcal efflux pump QacA. 2026.04.10.717755 Preprint at 10.64898/2026.04.10.717755 (2026).

32. Martens, C. et al. Direct protein-lipid interactions shape the conformational landscape of secondary transporters. Nat. Commun. 9, 4151 (2018).

33. Jia, R. et al. Hydrogen-deuterium exchange mass spectrometry captures distinct dynamics upon substrate and inhibitor binding to a transporter. Nat. Commun. 11, 6162 (2020).

34. Hariharan, P. et al. Allosteric effects of the coupling cation in melibiose transporter MelB. eLife https://elifesciences.org/articles/108335/peer-reviews (2026) doi:10.7554/eLife.108335.

35. Hariharan, P. et al. Mobile barrier mechanisms for Na+-coupled symport in an MFS sugar transporter. eLife https://elifesciences.org/articles/92462/figures (2024) doi:10.7554/eLife.92462.

36. Hariharan, P. et al. Mobile barrier mechanisms for Na+-coupled symport in an MFS sugar transporter. eLife 12, RP92462 (2024).

37. Hariharan, P. et al. Allosteric effects of the coupling cation in melibiose transporter MelB. eLife 14, RP108335 (2026).

38. Majumder, P. et al. Dissection of Protonation Sites for Antibacterial Recognition and Transport in QacA, a Multi-Drug Efflux Transporter. J. Mol. Biol. 431, 2163–2179 (2019).

39. Schaedler, T. A. & Van Veen, H. W. A flexible cation binding site in the multidrug major facilitator superfamily transporter LmrP is associated with variable proton coupling. FASEB J. 24, 3653–3661 (2010).

40. Nair, A. V. et al. Relocation of active site carboxylates in major facilitator superfamily multidrug transporter LmrP reveals plasticity in proton interactions. Sci. Rep. 6, 38052 (2016).

41. Discovery, C. et al. Chai-1: Decoding the molecular interactions of life. 2024.10.10.615955 Preprint at 10.1101/2024.10.10.615955 (2024).

42. Hageman, T. S. & Weis, D. D. Reliable Identification of Significant Differences in Differential Hydrogen Exchange-Mass Spectrometry Measurements Using a Hybrid Significance Testing Approach. Anal. Chem. 91, 8008–8016 (2019).

43. Lau, A. M., Claesen, J., Hansen, K. & Politis, A. Deuteros 2.0: peptide-level significance testing of data from hydrogen deuterium exchange mass spectrometry. Bioinformatics 37, 270–272 (2021).

44. Cecchetti, C. et al. A novel high-throughput screen for identifying lipids that stabilise membrane proteins in detergent based solution. PLOS ONE 16, e0254118 (2021).

45. Lacabanne, D. et al. Solid-State NMR Reveals Asymmetric ATP Hydrolysis in the Multidrug ABC Transporter BmrA. J. Am. Chem. Soc. 144, 12431–12442 (2022).

46. Selvasingh, J. A. et al. Dark nanodiscs for evaluating membrane protein thermostability by differential scanning fluorimetry. Biophys. J. 123, 68–79 (2024).

47. Kotov, V. et al. In-depth interrogation of protein thermal unfolding data with MoltenProt. Protein Sci. Publ. Protein Soc. 30, 201–217 (2021).

48. Martens, C. et al. Lipids modulate the conformational dynamics of a secondary multidrug transporter. Nat. Struct. Mol. Biol. 23, 744–751 (2016).

49. Maklad, H. R. et al. Antibiotic-Specific Conformational Landscapes of a Multidrug Transporter. 2025.05.17.654648 Preprint at 10.1101/2025.05.17.654648 (2026).

50. Quistgaard, E. M., Löw, C., Guettou, F. & Nordlund, P. Understanding transport by the major facilitator superfamily (MFS): structures pave the way. Nat. Rev. Mol. Cell Biol. 17, 123–132 (2016).

51. Jiang, D. et al. Structure of the YajR transporter suggests a transport mechanism based on the conserved motif A. Proc. Natl. Acad. Sci. 110, 14664–14669 (2013).

52. Mazurkiewicz, P., Driessen, A. J. M. & Konings, W. N. Energetics of wild-type and mutant multidrug resistance secondary transporter LmrP of *Lactococcus lactis*. Biochim. Biophys. Acta BBA - Bioenerg. 1658, 252–261 (2004).

53. Mazurkiewicz, P., Konings, W. N. & Poelarends, G. J. Acidic Residues in the Lactococcal Multidrug Efflux Pump LmrP Play Critical Roles in Transport of Lipophilic Cationic Compounds *. J. Biol. Chem. 277, 26081–26088 (2002).

54. Bapna, A. et al. Two Proton Translocation Pathways in a Secondary Active Multidrug Transporter. J. Mol. Microbiol. Biotechnol. 12, 197–209 (2007).

55. Kay, L. E. Protein dynamics from NMR. Nat. Struct. Biol. 5, 513–517 (1998).

56. Sekhar, A. & Kay, L. E. An NMR View of Protein Dynamics in Health and Disease. Annu. Rev. Biophys. 48, 297–319 (2019).

57. Dynamics of the Flexible Loop of Triose-Phosphate Isomerase: The Loop Motion Is Not Ligand Gated. ACS Publications https://pubs.acs.org/doi/pdf/10.1021/bi00026a012 (1995).

58. Eisenmesser, E. Z., Bosco, D. A., Akke, M. & Kern, D. Enzyme Dynamics During Catalysis. Science 295, 1520–1523 (2002).

59. Boehr, D. D., Dyson, H. J. & Wright, P. E. An NMR Perspective on Enzyme Dynamics. Chem. Rev. 106, 3055–3079 (2006).

60. Sanders, C. R. & Sönnichsen, F. Solution NMR of membrane proteins: practice and challenges. Magn. Reson. Chem. 44, S24–S40 (2006).

61. Mandala, V. S., Williams, J. K. & Hong, M. Structure and Dynamics of Membrane Proteins from Solid-State NMR. Annu. Rev. Biophys. 47, 201–222 (2018).

62. Oxenoid, K. & Chou, J. J. The present and future of solution NMR in investigating the structure and dynamics of channels and transporters. Curr. Opin. Struct. Biol. 23, 547–554 (2013).

63. Carrillo, V. H. P. et al. Bidirectional communication between nucleotide and substrate binding sites in a type IV multidrug ABC transporter. Nat. Commun. 16, 9921 (2025).

64. Aydin, D. et al. Molecular mechanism of exchange coupling in CLC chloride/proton antiporters. Nat. Commun. 17, 1342 (2026).

65. Coppieters ‘t Wallant, K. & Martens, C. Hydrogen-deuterium exchange coupled to mass spectrometry: A multifaceted tool to decipher the molecular mechanism of transporters. Biochimie 205, 95–101 (2023).

66. Giladi, M. & Khananshvili, D. Hydrogen-Deuterium Exchange Mass-Spectrometry of Secondary Active Transporters: From Structural Dynamics to Molecular Mechanisms. Front. Pharmacol. 11, (2020).

67. Möller, I. R. et al. Conformational dynamics of the human serotonin transporter during substrate and drug binding. Nat. Commun. 10, 1687 (2019).

68. Reddy, K. D., Ciftci, D., Scopelliti, A. J. & Boudker, O. The archaeal glutamate transporter homologue GltPh shows heterogeneous substrate binding. J. Gen. Physiol. 154, e202213131 (2022).

69. Adhikary, S., et al. Conformational dynamics of a neurotransmitter:sodium symporter in a lipid bilayer. Proc. Natl. Acad. Sci. 114, E1786–E1795 (2017).

70. Malinauskaite, L. et al. A mechanism for intracellular release of Na+ by neurotransmitter/sodium symporters. Nat. Struct. Mol. Biol. 21, 1006–1012 (2014).

71. Hou, C., Zhao, H. & Shen, Y. Protein language models trained on biophysical dynamics inform mutation effects. Proc. Natl. Acad. Sci. 123, e2530466123 (2026).

72. Gavalda-Garcia, J., Dixit, B., Díaz, A., Ghysels, A. & Vranken, W. Gradations in protein dynamics captured by experimental NMR are not well represented by AlphaFold2 models and other computational metrics. J. Mol. Biol. 437, 168900 (2025).

73. Sledzieski, S. & Hanson, S. RocketSHP: Ultra-fast Proteome-scale Prediction of Protein Dynamics. 2025.06.12.659353 Preprint at 10.1101/2025.06.12.659353 (2025).

74. High-Frequency Transformation, by Electroporation, of Lactococcus lactis subsp. cremoris Grown with Glycine in Osmotically Stabilized Media. https://journals.asm.org/doi/epdf/10.1128/aem.55.12.3119-3123.1989?src=getftr&utm_source=acs&getft_integrator=acsdoi:10.1128/aem.55.12.3119-3123.1989.

75. del Alamo, D., Govaerts, C. & Mchaourab, H. S. AlphaFold2 predicts the inward-facing conformation of the multidrug transporter LmrP. Proteins Struct. Funct. Bioinforma. 89, 1226–1228 (2021).

