## Supplemental Figures for "Local conformational plasticity underlies ligand recognition and proton coupling in MFS multidrug transporters"

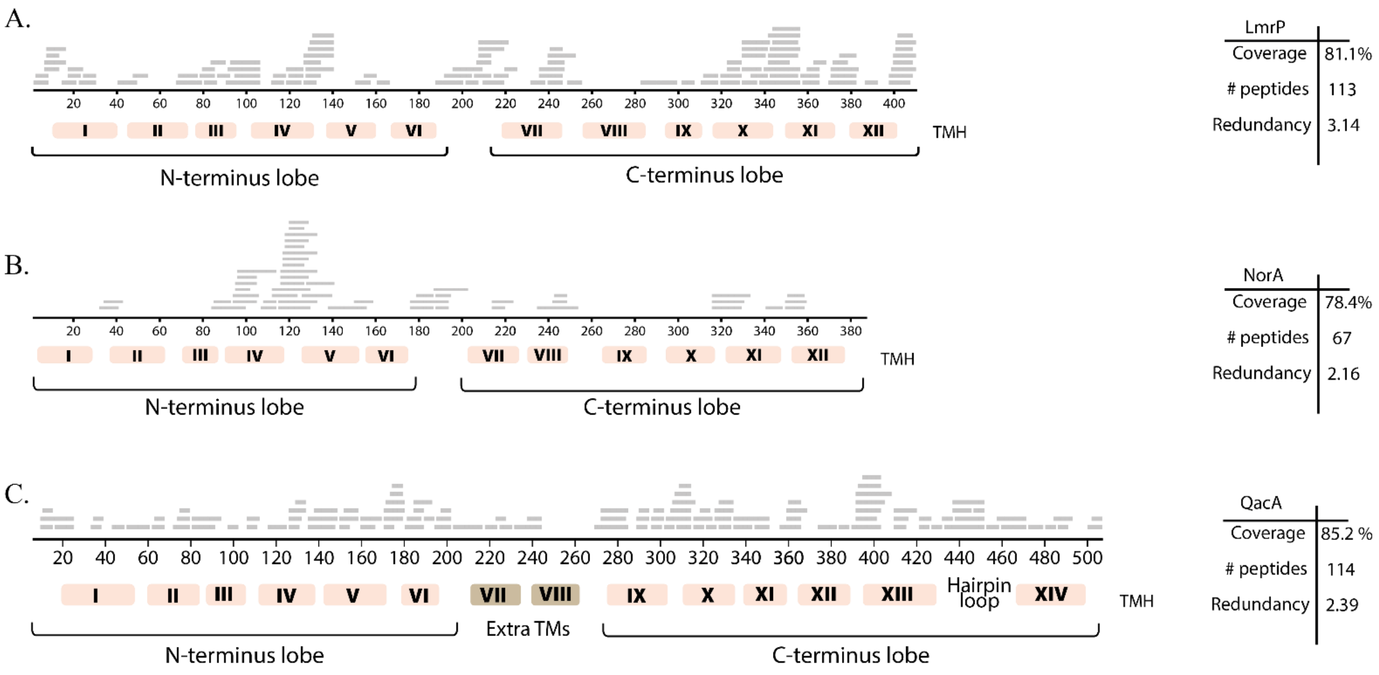


**Supplementary Figure 1 :** Coverage maps of pepsin proteolyzed peptides of LmrP (**A**), NorA (**B**) and QacA (**C**).

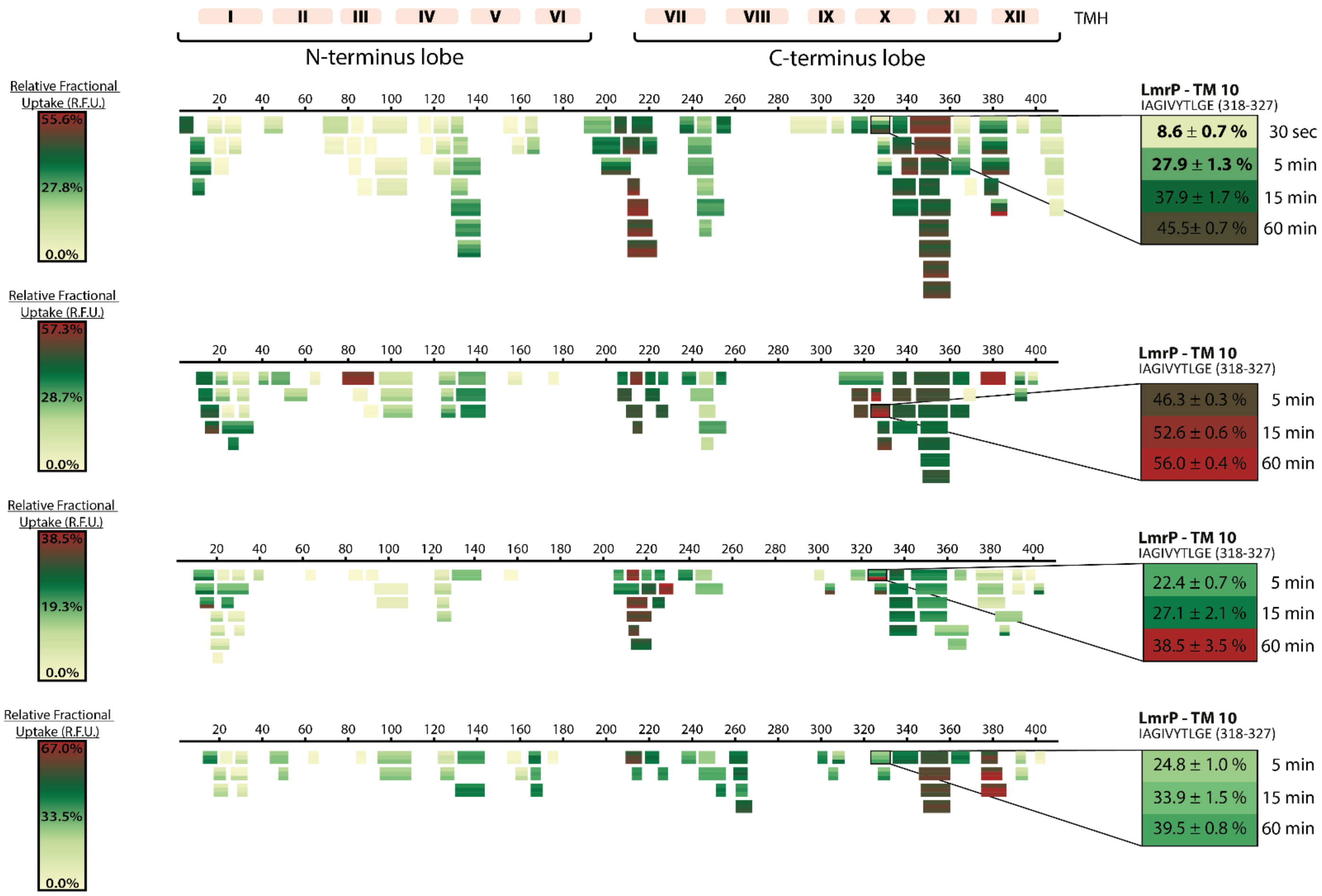


**Supplementary Figure 2. Peptide coverage and deuterium uptake maps for four biological replicates of LmrP.** Coverage maps show the relative fractional uptake (R.F.U.) measured for each identified peptide at each monitored labelling time point. Peptides are positioned according to their location in the LmrP sequence and transmembrane topology. Temporal exchange data are displayed separately for each peptide. The four biological replicates are shown independently, with replicate-specific R.F.U colour scales to account for differences in overall exchange range. The peptide-level views on the right illustrate the exchange behaviour for a representative TM10 peptide across replicates.

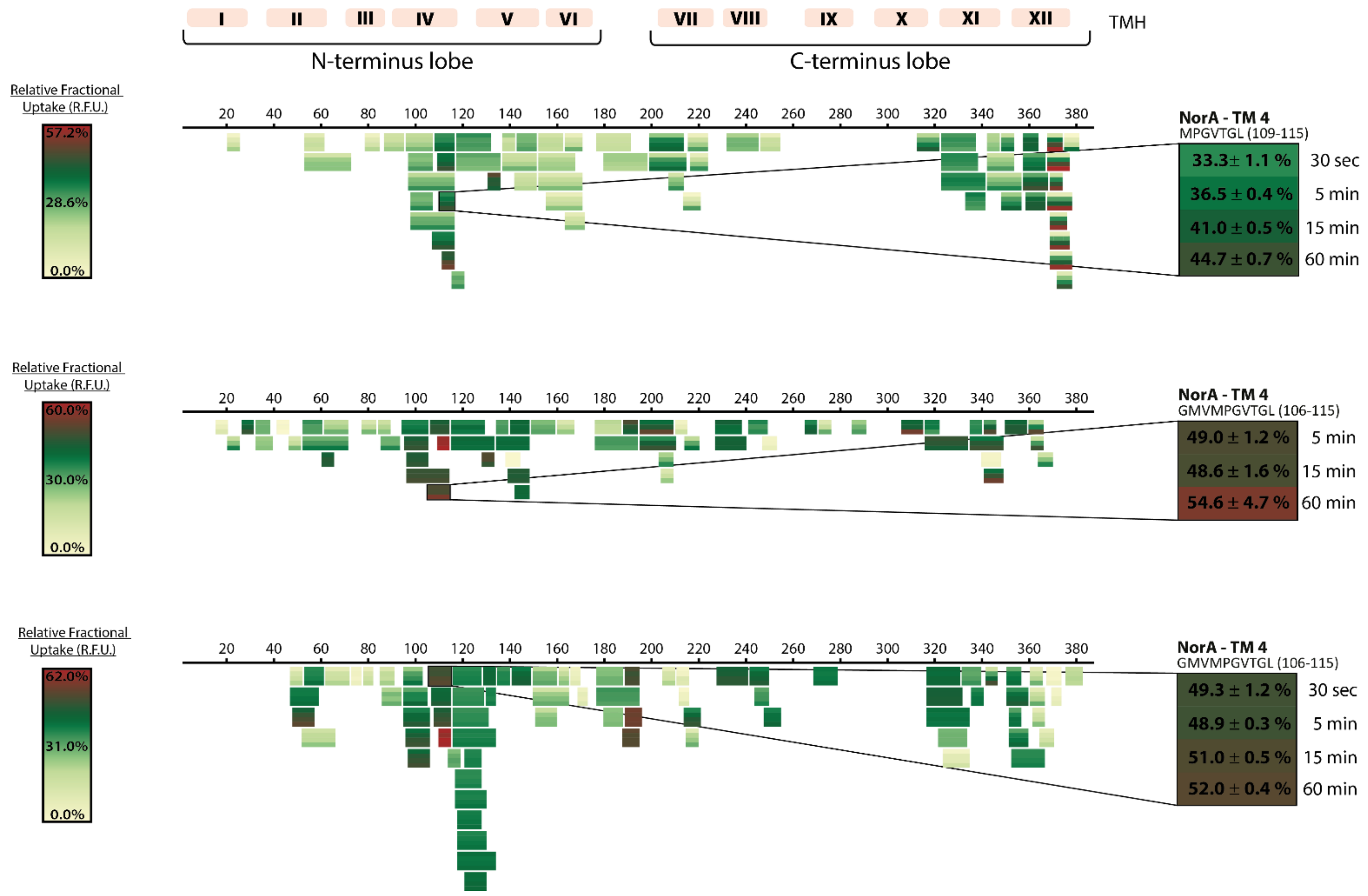


**Supplementary Figure 3. Peptide coverage and deuterium uptake maps for three biological replicates of NorA.** Coverage maps show the relative fractional uptake (R.F.U.) measured for each identified peptide at each monitored labelling time point. Peptides are positioned according to their location in the NorA sequence and transmembrane topology. Temporal exchange data are displayed separately for each peptide. The three biological replicates are shown independently, with replicate-specific R.F.U colour scales to account for differences in overall exchange range. The peptide-level views on the right illustrate the exchange behaviour for a representative TM4 peptide across replicates.

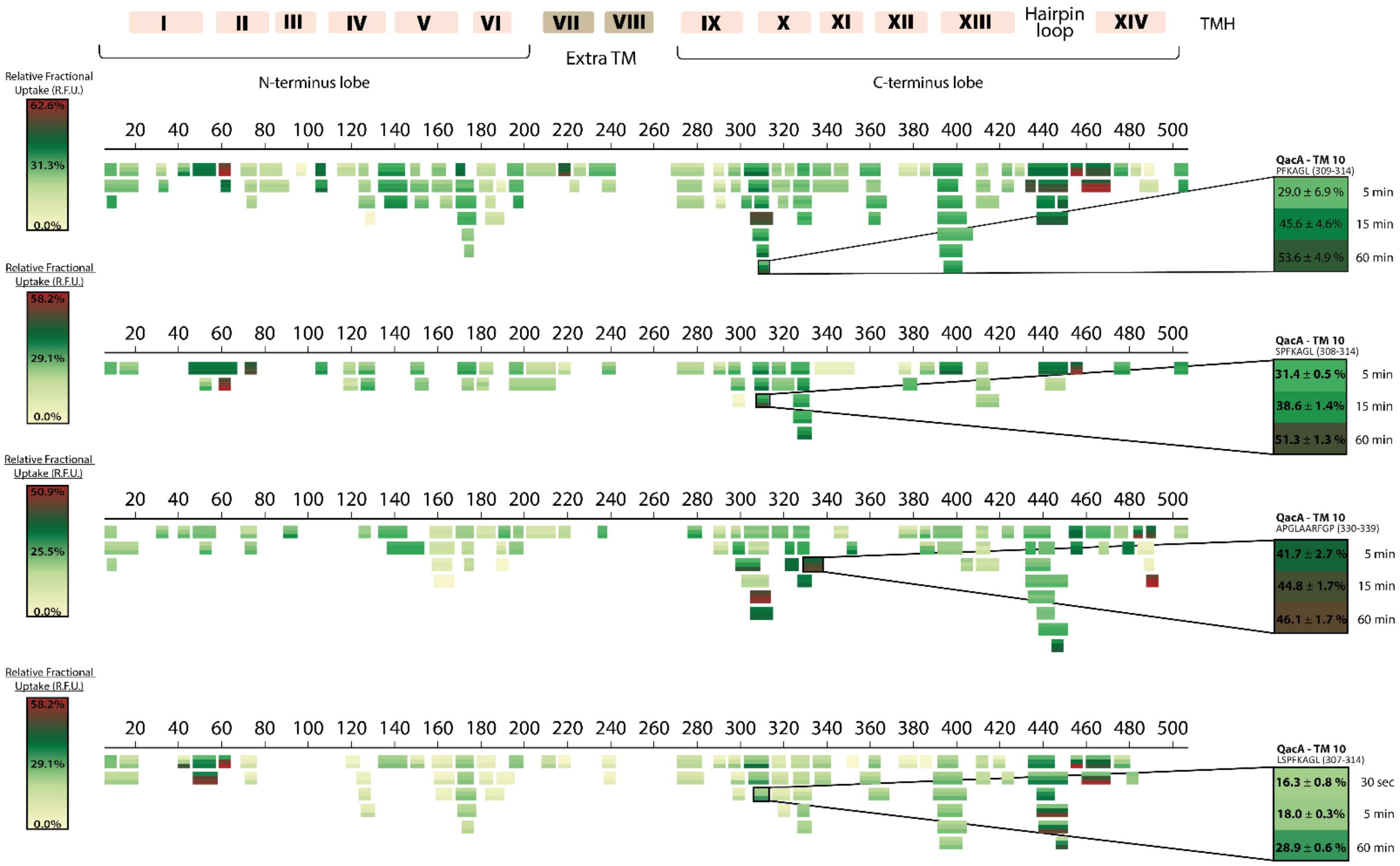


**Supplementary Figure 4. Peptide coverage and deuterium uptake maps for three biological replicates of QacA.** Coverage maps show the relative fractional uptake (R.F.U.) measured for each identified peptide at each monitored labelling time point. Peptides are positioned according to their location in the QacA sequence and transmembrane topology. Temporal exchange data are displayed separately for each peptide. The biological replicates are shown independently, with replicate-specific R.F.U colour scales to account for differences in overall exchange range. The peptide-level views on the right illustrate the exchange behaviour for a representative TM10 peptide across replicates.

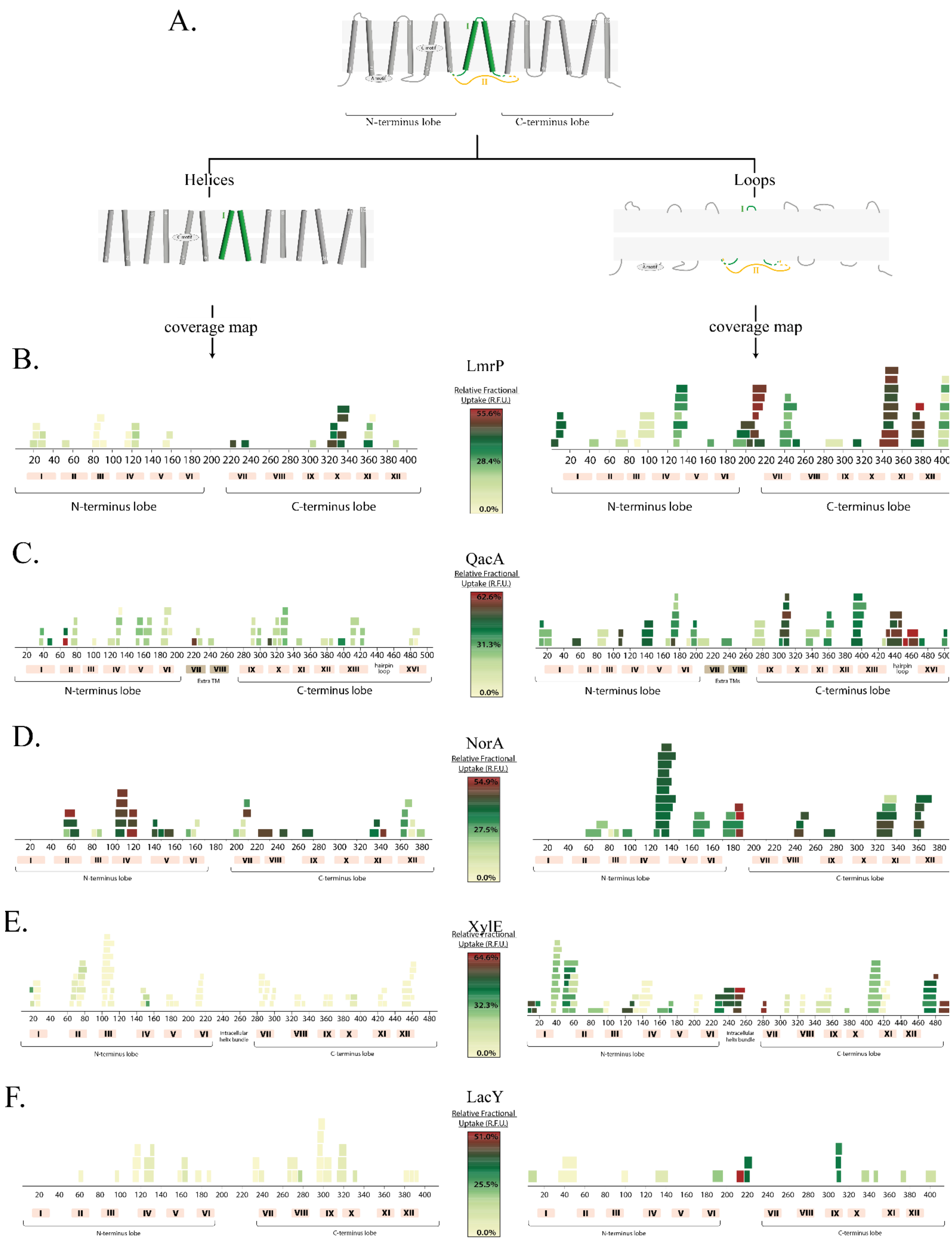


**Supplementary Figure 5. Definition of transmembranous and extramembranous peptide subsets across MFS transporters.** (A) Workflow used to partition the peptide-level datasets into helix and loop/extramembrane subsets. Peptide assignment was guided by available structural information, including AlphaFold-predicted models and experimentally determined structures, to distinguish segments mapping to transmembrane helices from those located in loop or non-helical regions. (B-F) Application of this classification to MFS transporters that have been subjected to HDX-MS analysis: LmrP, QacA, NorA, XylE and LacY, shown from top to bottom. The data for XylE and LacY was downloaded from ProteomeXchange Consortium PRIDE (dataset identifier PXD011060). This subdivision was used to compare exchange behaviour between transmembrane helical regions and loop/extramembrane regions across different members of the MFS fold.

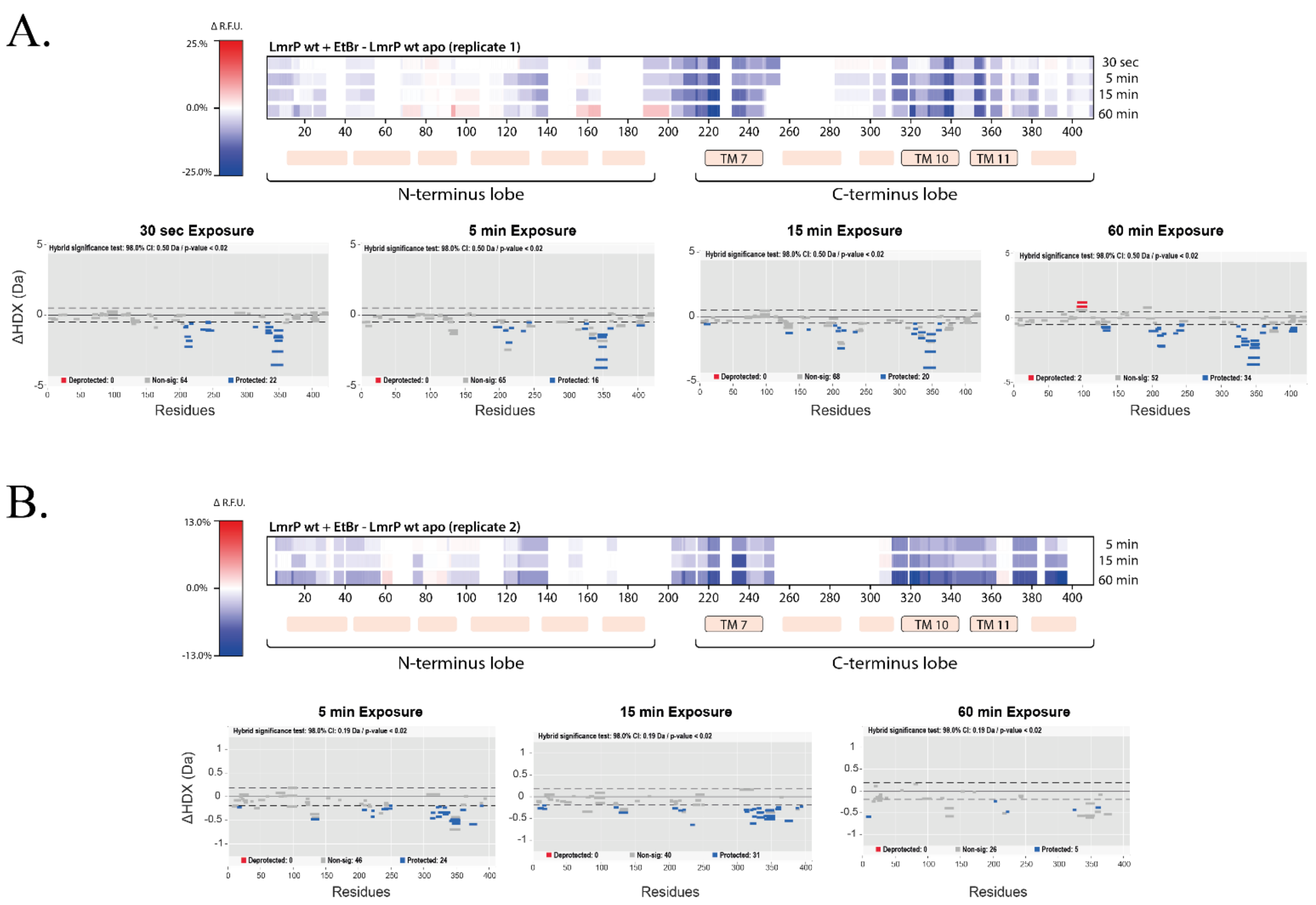


**Supplementary Figure 6**: **Effect of ethidium bromide on LmrP dynamics across two independent HDX-MS replicates.** ΔRFU heatmaps show the change in deuterium uptake induced by ethidium bromide (EtBr) binding in two independent replicates(A,B). Values correspond to the difference between the EtBr-bound and apo states, mapped onto the LmrP sequence and transmembrane-helix annotation. Each replicate is displayed using an independently normalized RFU scale. The corresponding Woods plots for each time point are shown below each heatmap. Peptides showing a statistically significant increase or decrease in uptake are shown in red and blue, respectively, non-significant peptides are shown in grey. Statistical significance was assessed in Deuteros using $\alpha = 0.02$ and hybrid analysis[1].

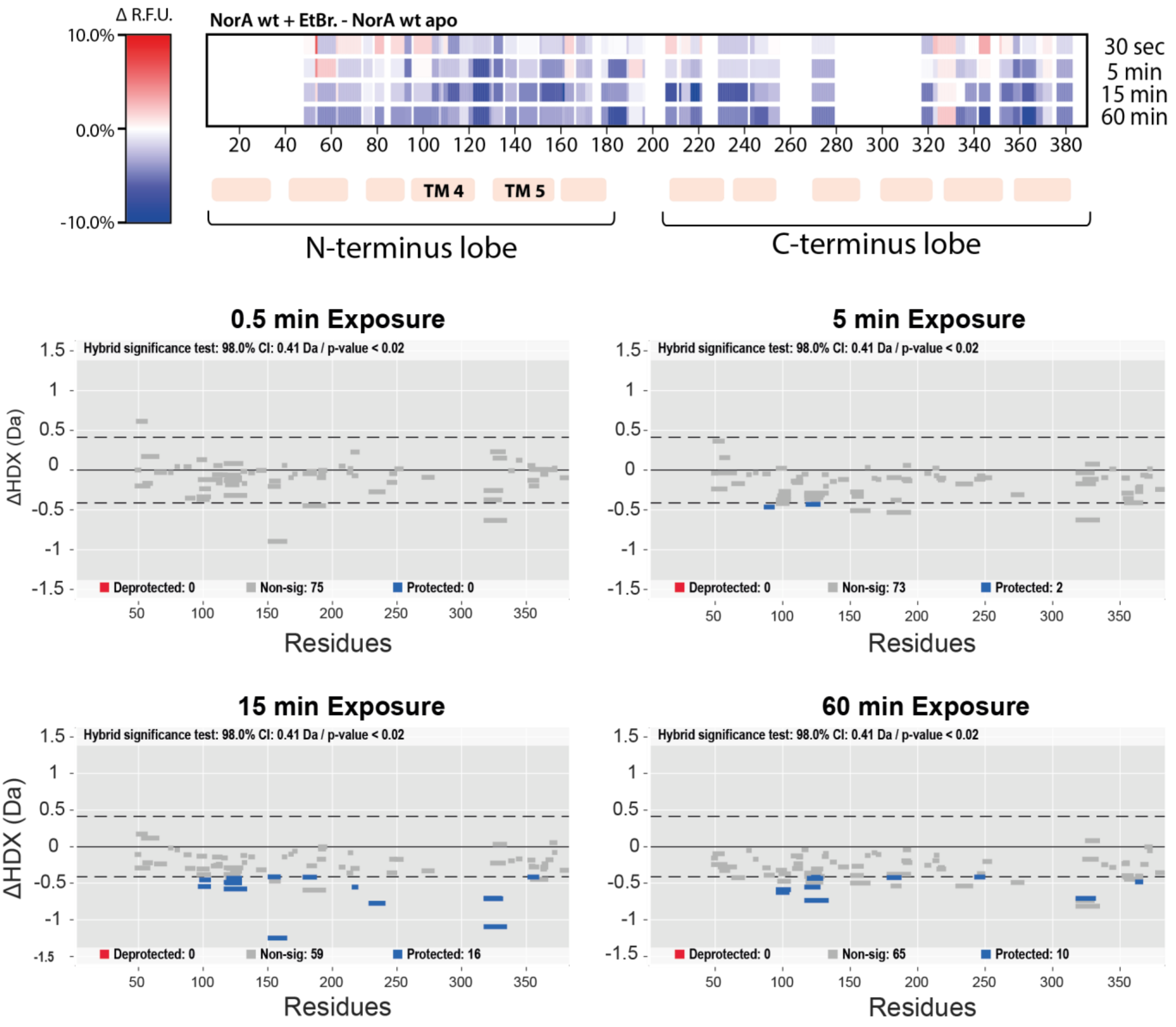


**Supplementary Figure 7**: **Effect of ethidium bromide on NorA dynamics.** ΔRFU heatmaps show the change in deuterium uptake induced by ethidium bromide (EtBr) binding. Values correspond to the difference between the EtBr-bound and apo states, mapped onto the NorA sequence and transmembrane-helix annotation. The corresponding Woods plots for each time point is shown below the heatmap. Peptides showing a statistically significant increase or decrease in uptake are shown in red and blue, respectively, non-significant peptides are shown in grey. Statistical significance was assessed in Deuteros using α = 0.02 and hybrid analysis[1].

A.

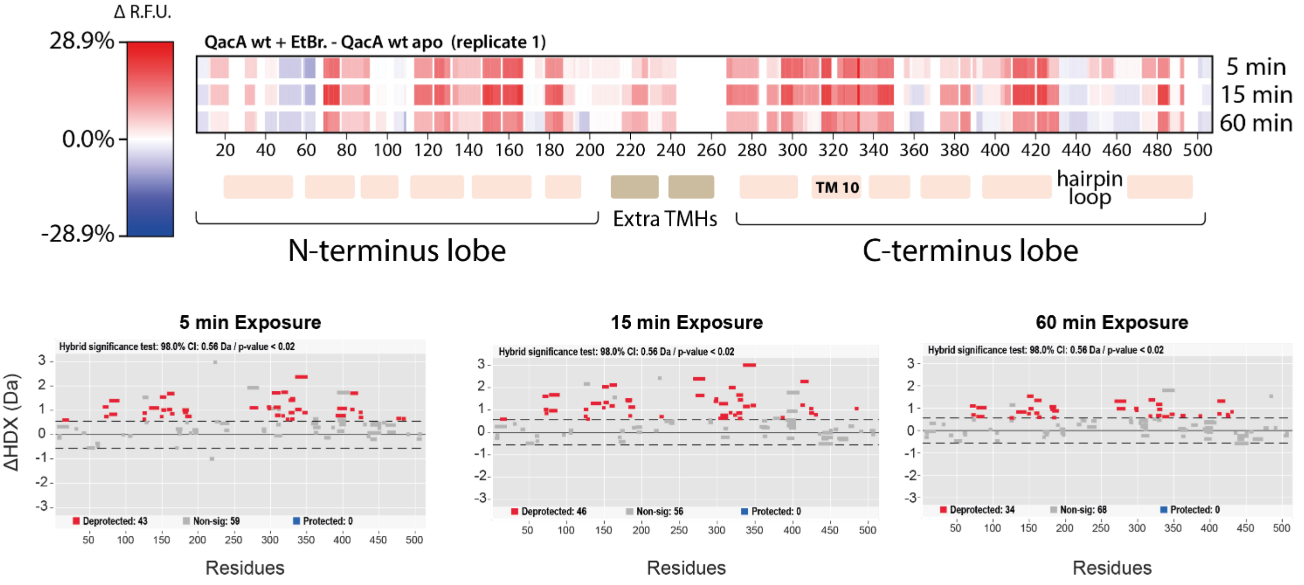


B.

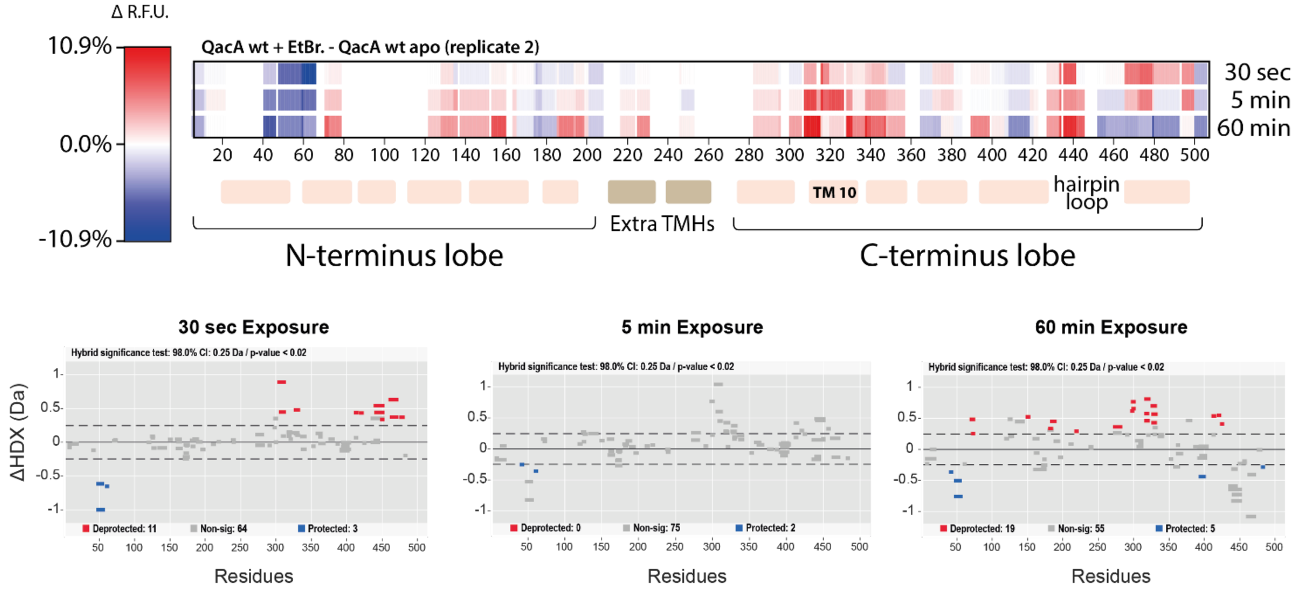


**Supplementary Figure 8: Effect of ethidium bromide on QacA dynamics across two independent HDX-MS replicates.** ΔRFU heatmaps show the change in deuterium uptake induced by ethidium bromide (EtBr) binding in two independent replicates(A,B). Values correspond to the difference between the EtBr-bound and apo states, mapped onto the QacA sequence and transmembrane-helix annotation. Each replicate is displayed using an independently normalized RFU scale. The corresponding Woods plots for each time point are shown below each heatmap. Peptides showing a statistically significant increase or decrease in uptake are shown in red and blue, respectively, non-significant peptides are shown in grey. Statistical significance was assessed in Deuteros using α = 0.02 and hybrid analysis[1].

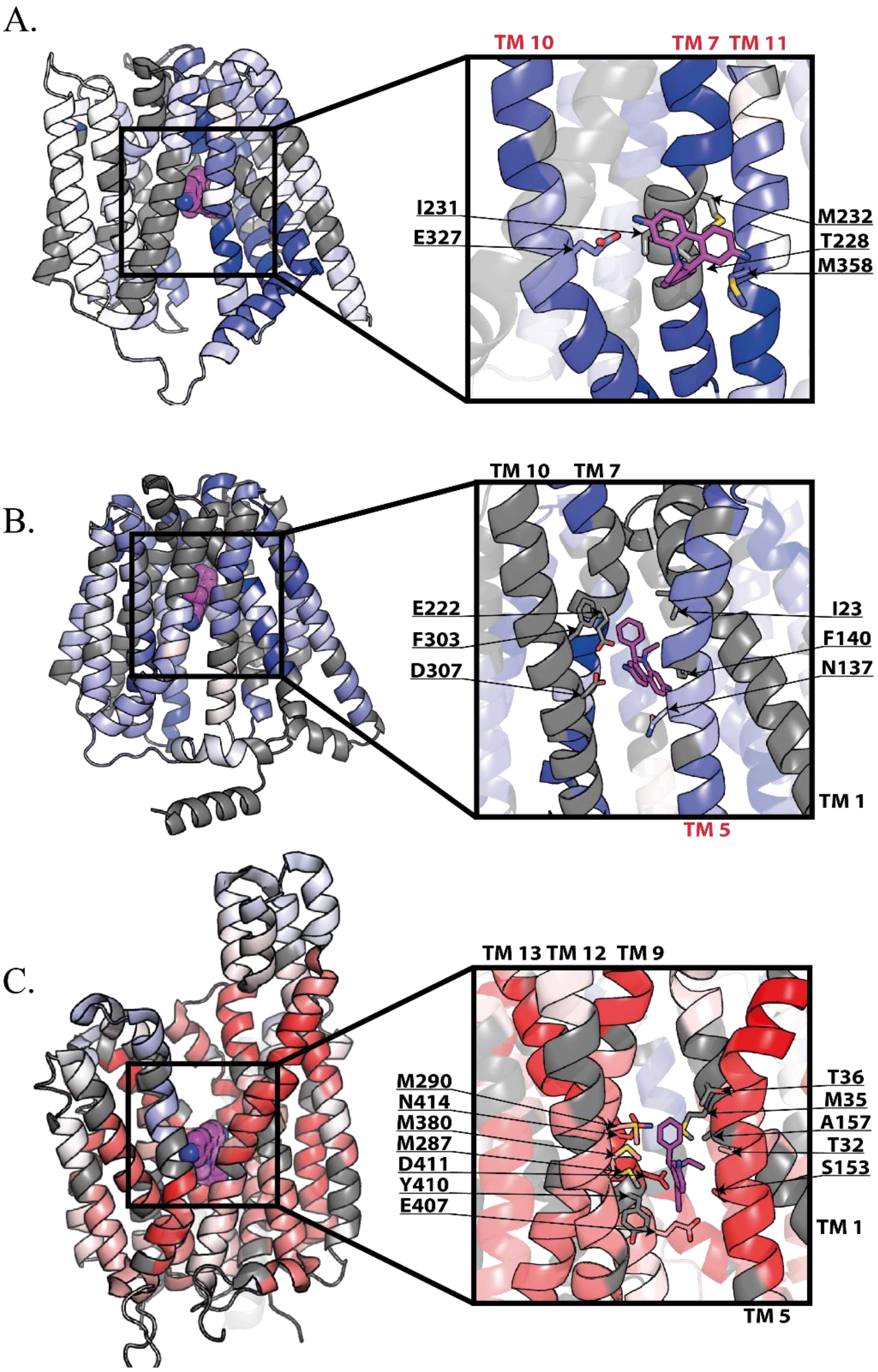


**Supplementary Figure 9. Structural environment of ethidium in the binding pocket of LmrP, NorA and QacA.** Full-transporter views and corresponding close-up views of the ethidium bromide (EtBr)-binding pocket are shown for LmrP (A), NorA (B) and QacA (C). LmrP and NorA were modelled with Chai-1[2], whereas QacA is shown from the experimental structure, PDB 29PX. EtBr is shown as magenta sticks, and residues located within 4 Å of the ligand are shown as sticks in the enlarged views. Relevant transmembrane helices are indicated. The protein cartoons are coloured according to the ethidium-induced HDX-MS response, highlighting the local structural context of ligand binding relative to regions showing altered deuterium uptake.

A.

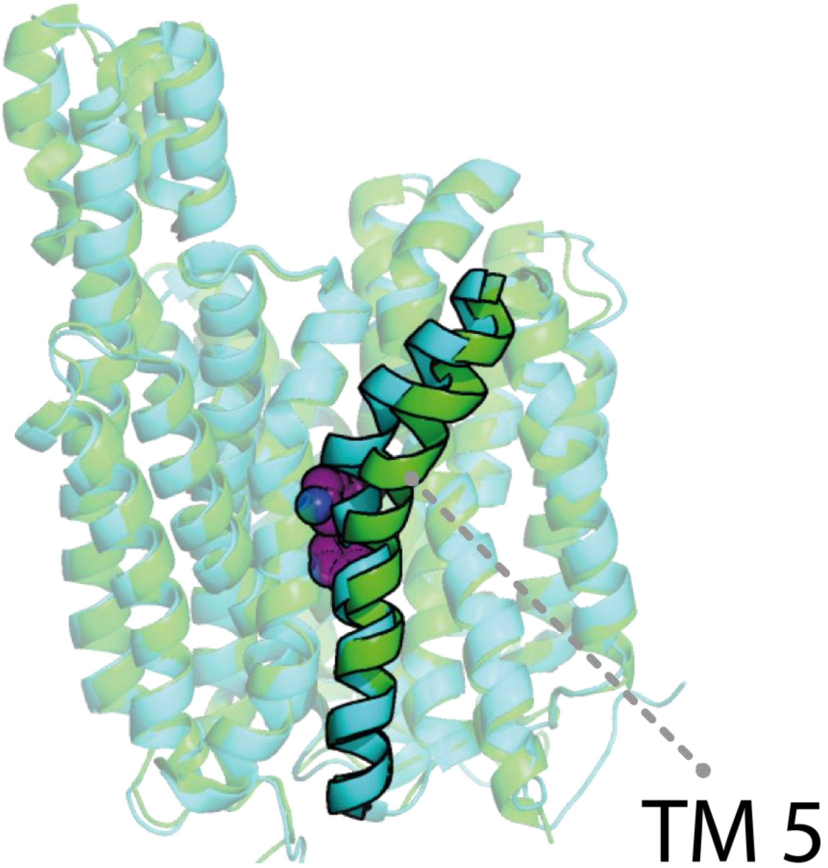


**Supplementary Figure 10. Ethidium bromide binding is associated with a displacement of QacA TMH5.** Structural overlay of QacA in the apo state (cyan; PDB: 7Y58) and in the ethidium bromide-bound state (green; PDB: 29PX). The overall transporter fold is largely conserved between the two structures, whereas TMH5 adopts a shifted position in the ligand-bound state. This movement, highlighted in the overlay, enlarges and remodels the central binding pocket to accommodate ethidium bromide, shown in magenta.

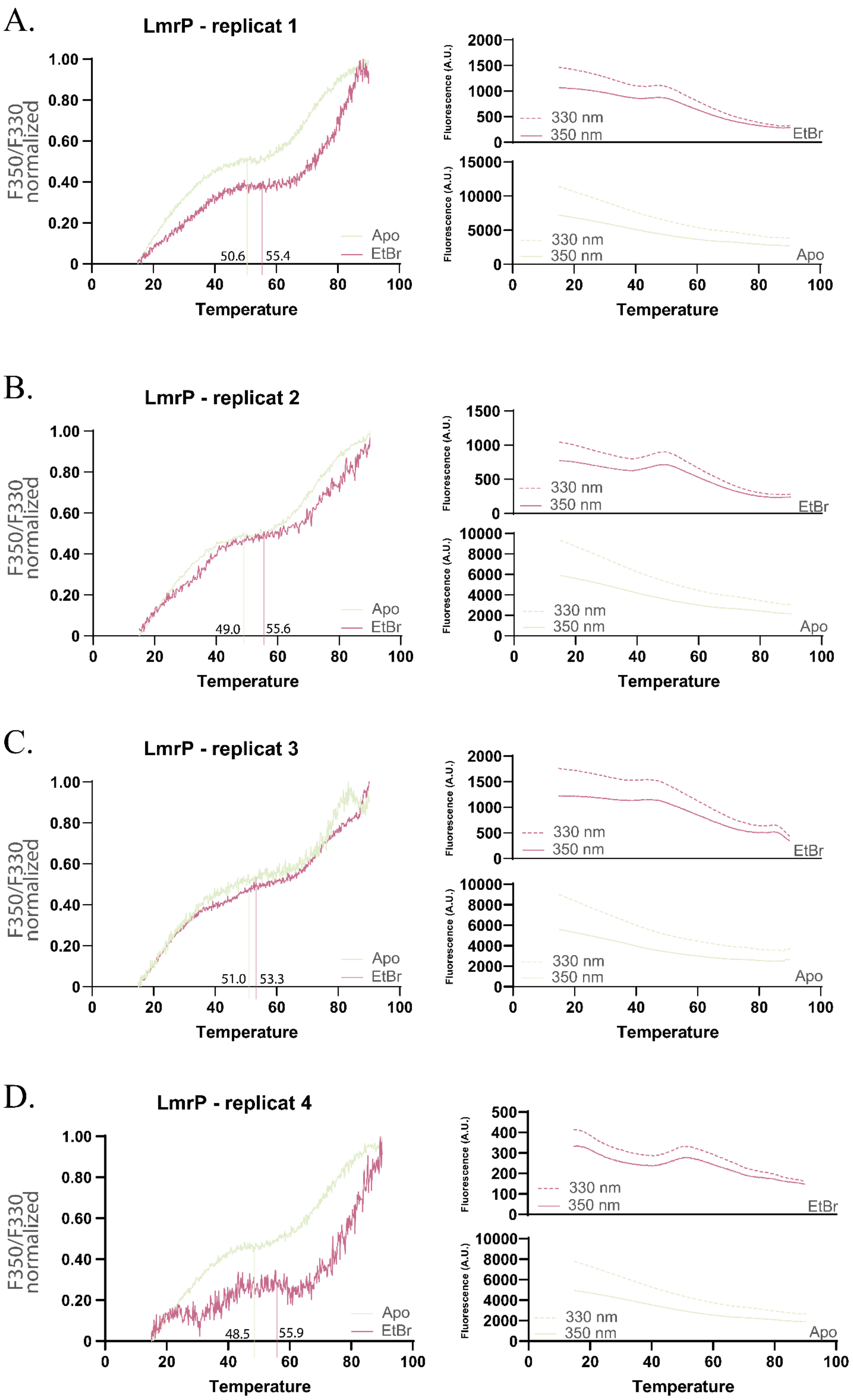
A.
LmrP - replicat 1
F350/F330 normalized
1.00
0.80
0.60
0.40
0.20
0
50.6
55.4
Apo
EtBr
0 20 40 60 80 100
Temperature
Fluorescence (A.U.)
2000 1500 1000 500 0
330 nm
350 nm
EtBr
15000 10000 5000 0
330 nm
350 nm
Apo
0 20 40 60 80 100
Temperature
B.
LmrP - replicat 2
F350/F330 normalized
49.0
55.6
Apo
EtBr
Temperature
1500 1000 500 0
330 nm
350 nm
EtBr
10000 8000 6000 4000 2000 0
330 nm
350 nm
Apo
Temperature
C.
LmrP - replicat 3
F350/F330 normalized
51.0
53.3
Apo
EtBr
Temperature
2000 1500 1000 500 0
330 nm
350 nm
EtBr
10000 8000 6000 4000 2000 0
330 nm
350 nm
Apo
Temperature
D.
LmrP - replicat 4
F350/F330 normalized
48.5
55.9
Apo
EtBr
Temperature
500 400 300 200 100 0
330 nm
350 nm
EtBr
10000 8000 6000 4000 2000 0
330 nm
350 nm
Apo
Temperature

**Supplementary Figure 11** : **Reproducibility of the nanoDSF thermal unfolding profiles of LmrP in the presence of ethidium bromide.** nanoDSF analysis of LmrP in the apo state and in the presence of 125 μM ethidium bromide (EtBr) across four independent biological replicates. For each replicate, the left panel shows the normalized F350/F330 fluorescence ratio as a function of temperature, with vertical lines indicating the apparent transition temperature, $T_{app}$, determined using the tangent-extrapolation method described in the Methods. The right panel shows the corresponding raw fluorescence emission traces at 330 nm and 350 nm for apo LmrP and EtBr-bound LmrP. Across replicates, LmrP displays reproducible thermal unfolding profiles, with EtBr producing a consistent shift in the apparent transition while preserving the overall shape of the unfolding curve.

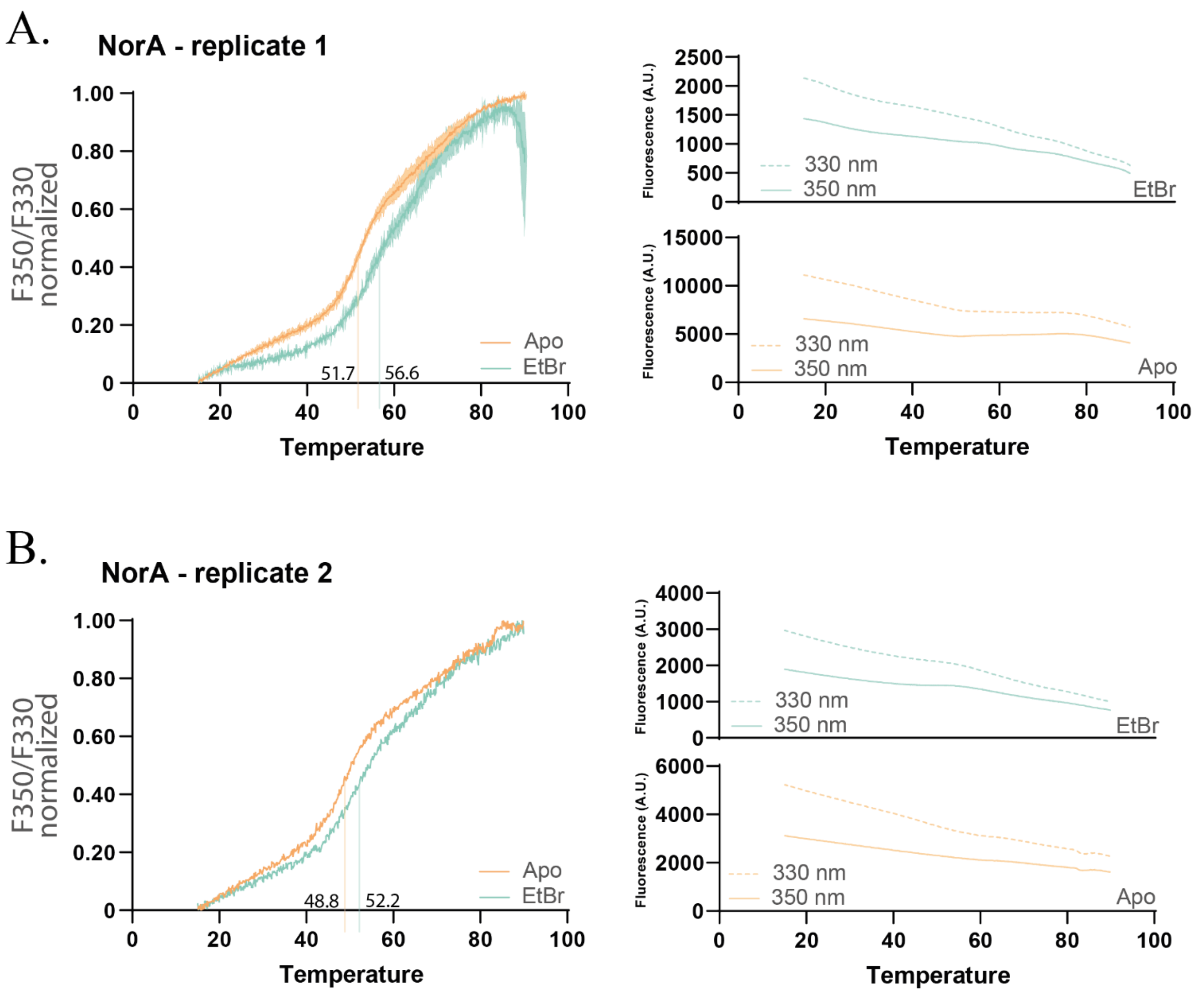


**Supplementary Figure 12 : Thermal stability analysis of NorA in the presence of ethidium bromide (EtBr).** nanoDSF analysis of NorA in the apo state and in the presence of 125 µM ethidium bromide (EtBr) across two independent biological replicates. For each replicate, the left panel shows the normalized F350/F330 fluorescence ratio as a function of temperature, with vertical lines indicating the apparent transition temperature, $T_{app}$, determined using the tangent-extrapolation method described in the Methods. The right panel shows the corresponding raw fluorescence emission traces at 330 nm and 350 nm for apo NorA and EtBr-bound NorA.

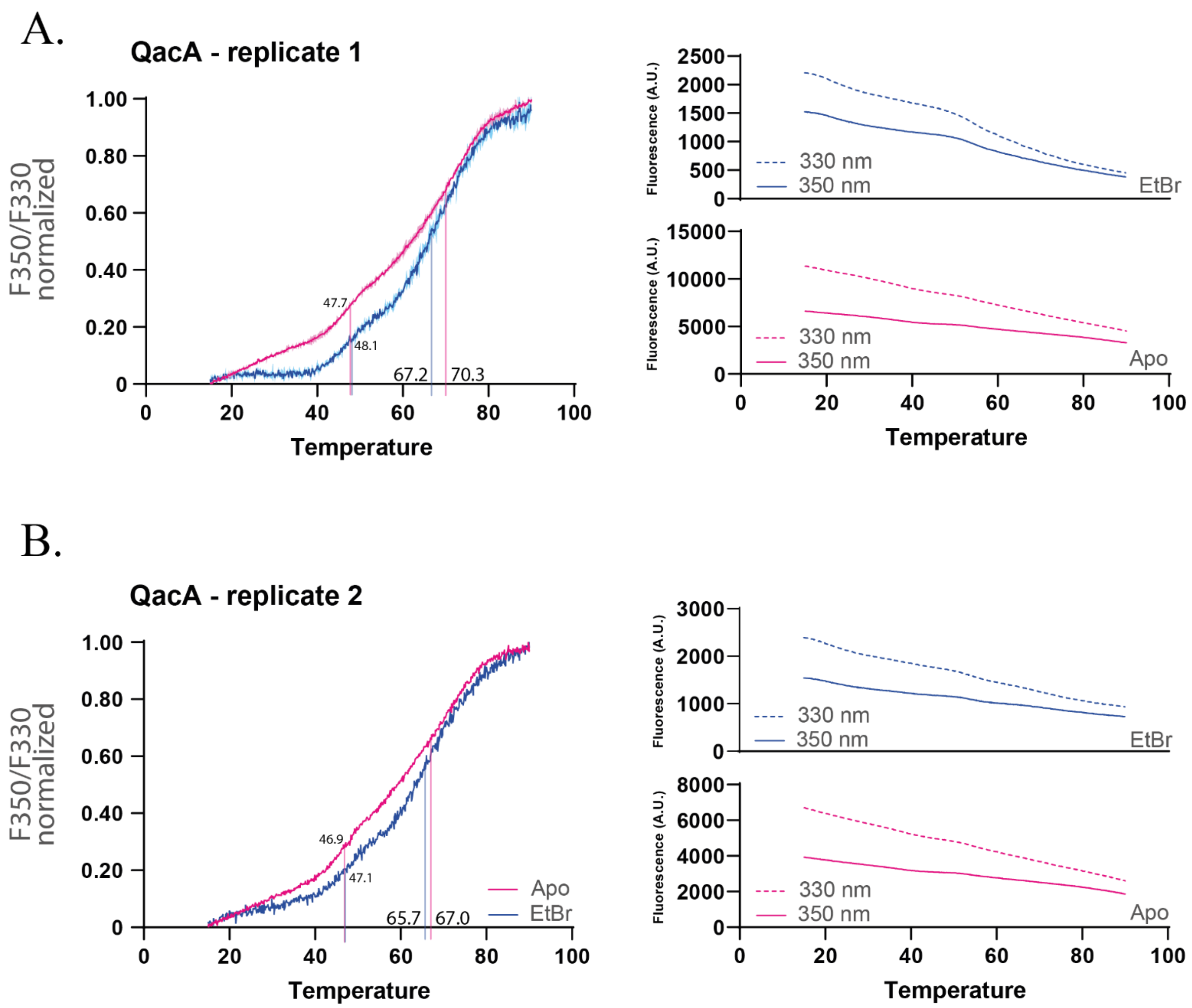


**Supplementary Figure 13 : Thermal stability analysis of QacA in the presence of ethidium bromide (EtBr).** nanoDSF analysis of QacA in the apo state and in the presence of 125 μM ethidium bromide (EtBr) across two independent biological replicates. For each replicate, the left panel shows the normalized F350/F330 fluorescence ratio as a function of temperature, with vertical lines indicating the apparent transition temperature, $T_{app}$, determined using the tangent-extrapolation method described in the Methods. The right panel shows the corresponding raw fluorescence emission traces at 330 nm and 350 nm for apo QacA and EtBr-bound QacA.

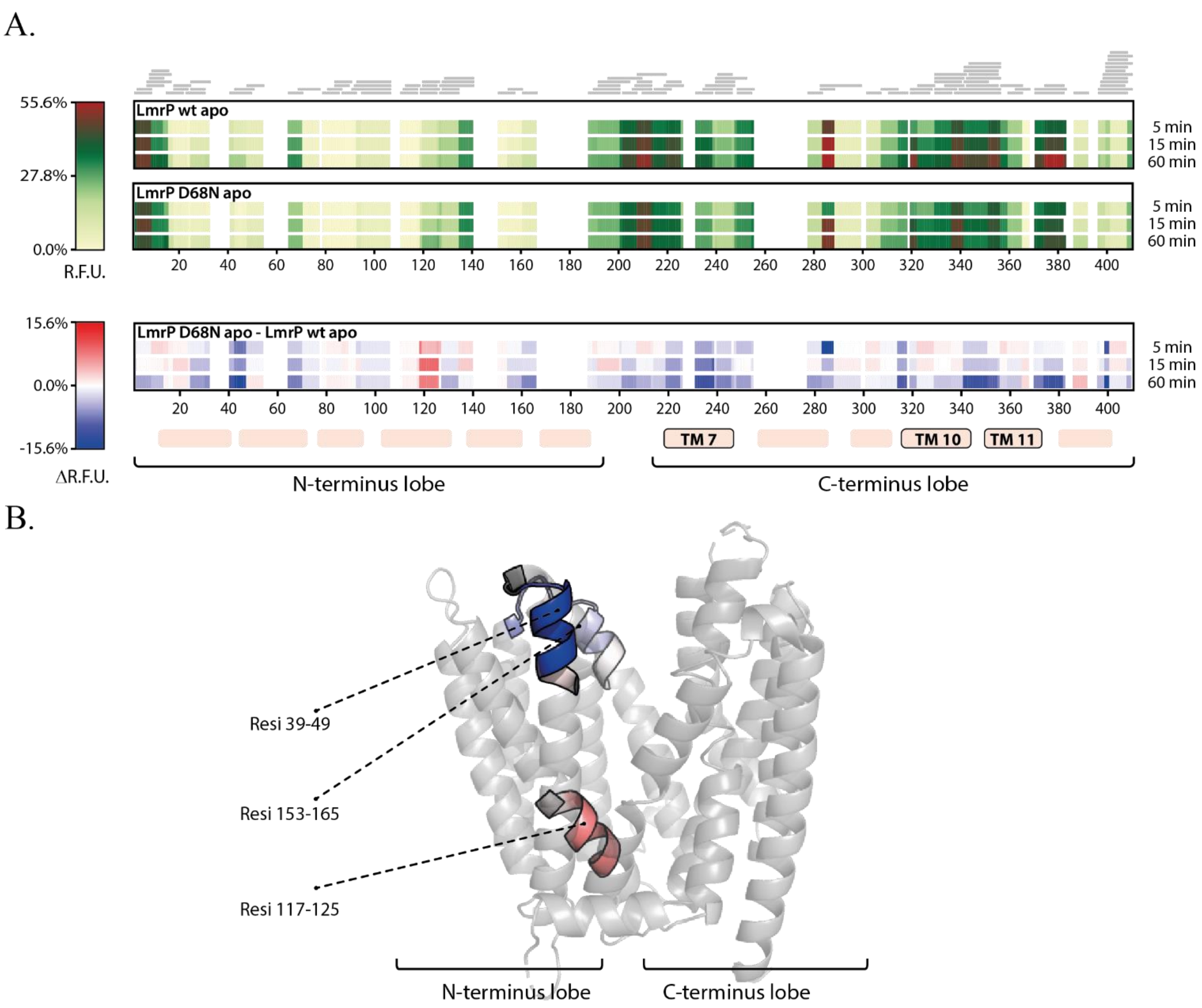


**Supplementary Figure 14 :The D68N mutation yields a ΔR.F.U pattern characteristic of a shift towards an inward facing state in the N-lobe of apo LmrP.** (A) Comparative HDX-MS heatmap analysis of apo wild-type LmrP and apo LmrP D68N. From top to bottom, heatmaps show deuterium uptake in wild-type LmrP, deuterium uptake in LmrP D68N, and the difference between the two states. Uptake values were displayed using a common scale normalized to the maximum uptake observed for apo wild-type LmrP (55.6%); the maximum uptake observed for apo LmrP D68N was approximately 50%. The differential heatmap highlights regions with altered exchange in the D68N mutant relative to the wild type. (B) Structural mapping of regions showing significant HDX-MS differences across multiple labelling time points. Peptides spanning residues 39–49 and 153–165 show protection in LmrP D68N, whereas residues 117–125 show increased exchange. These changes localize to the N-lobe and are consistent with the conformational rearrangements associated with the D68N-induced shift in the LmrP conformational equilibrium.

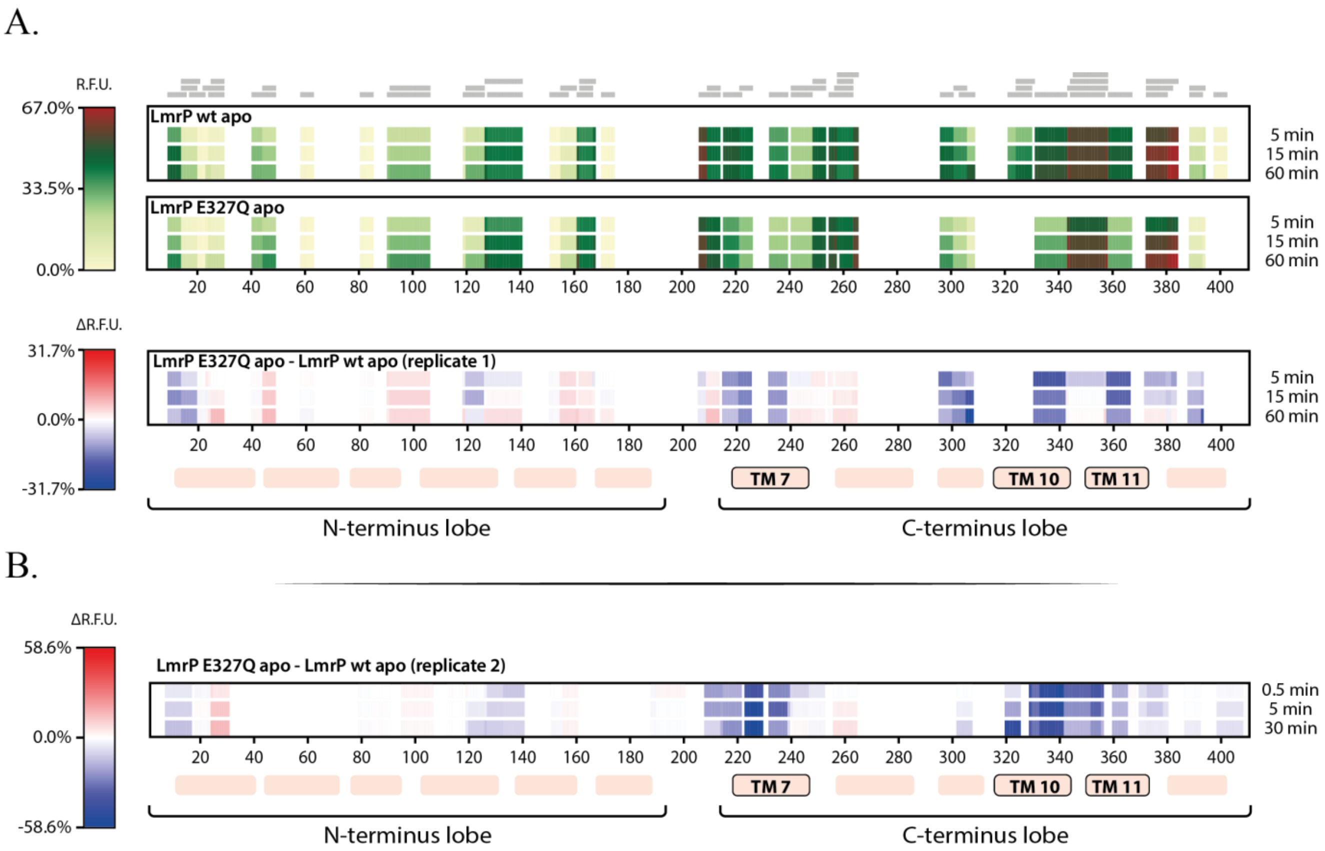


**Supplementary Figure 15 : Effect of the E327Q mutation on the local structural dynamics of apo LmrP.** (A) Comparative RFU heatmaps of apo LmrP WT and apo LmrP E327Q, aligned along the LmrP sequence. The upper heatmaps show the deuterium uptake profiles for WT and E327Q, respectively, while the lower heatmap shows the corresponding difference map between E327Q and WT. Positive values indicate increased exchange in E327Q relative to WT, whereas negative values indicate protection from exchange. Transmembrane helices are indicated below the heatmaps, with TM7, TM10 and TM11 highlighted (B) Independent biological replicate showing the same E327Q vs WT comparison.

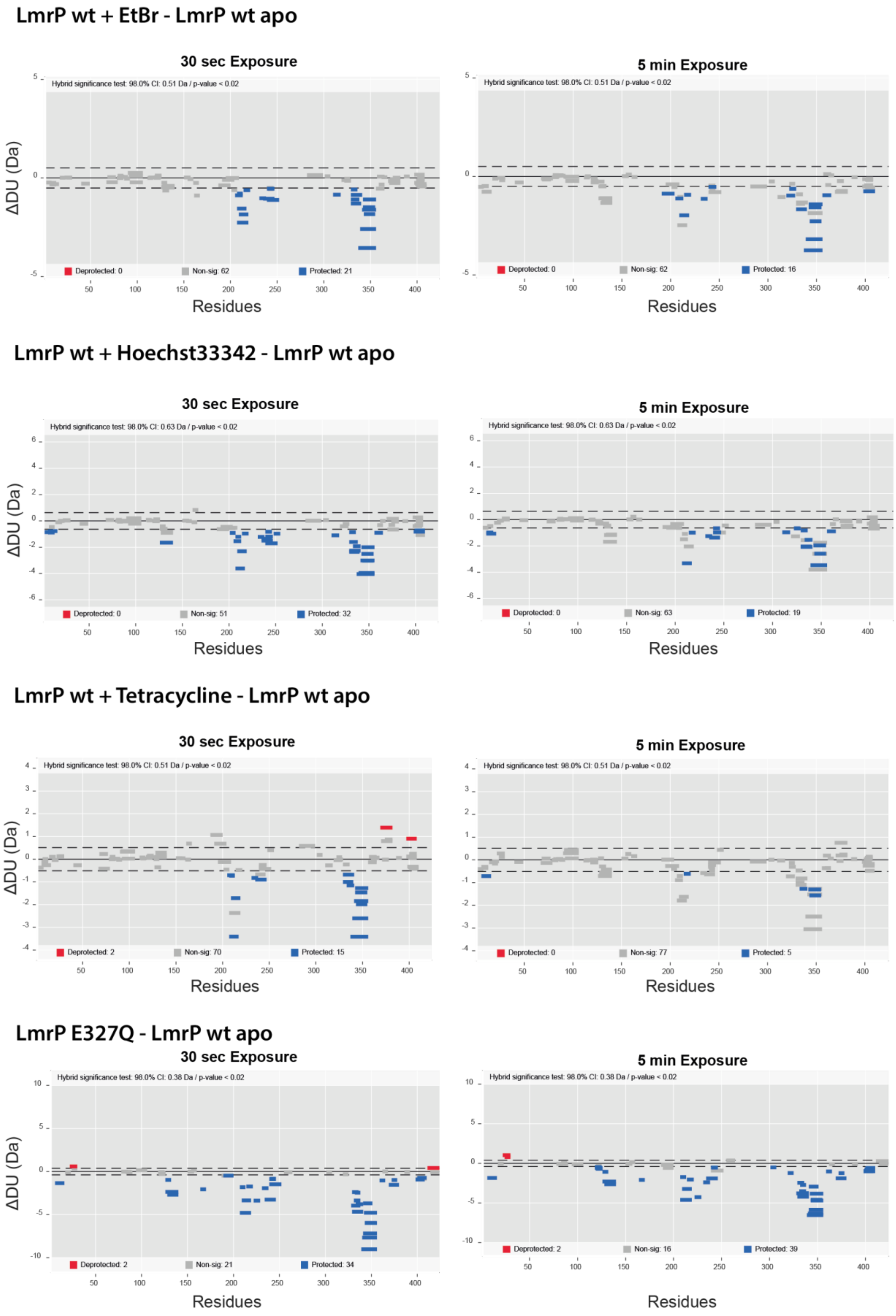


**Supplementary Figure 16. Woods plots of substrate and E327Q-induced changes in LmrP deuterium uptake.** Woods plots showing peptide-level differences in deuterium uptake between apo LmrP WT and LmrP WT in the presence of ethidium bromide, Hoechst 33342, or tetracycline, as well as between apo LmrP WT and apo LmrP E327Q. Comparisons are shown after 30 s and 5 min of deuterium labelling. Each horizontal bar represents an identified peptide mapped along the LmrP sequence, with the y-axis indicating the difference in deuterium uptake. Blue peptides show significant protection from exchange, red peptides show significant deprotection, and grey peptides show non-significant changes, as determined by the hybrid significance test (a=0.02%)[3]. Dashed lines indicate the significance thresholds. Hoechst 33342 and tetracycline both induce protection in the same flexible C-terminal helical region affected by ethidium binding and by the E327Q mutation, including TMHs 7, 10 and 11. The Woods plots were generated with Deuteros 2.0[1]

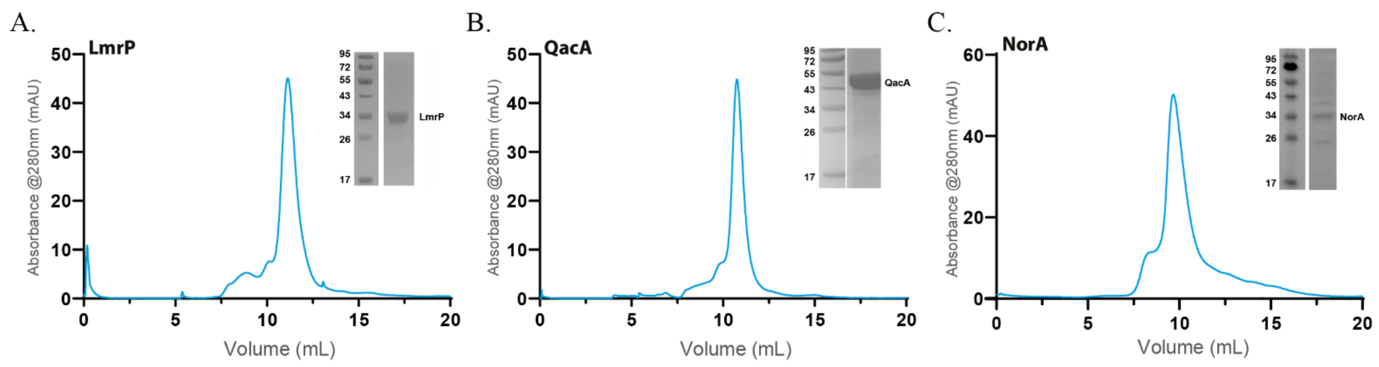


**Supplementary Figure 17. Purification profiles of LmrP, QacA and NorA.** Representative size-exclusion chromatography profiles of purified LmrP, QacA and NorA in detergent micelles, monitored by absorbance at 280 nm. The main elution peak corresponding to each transporter is shown. Insets show the corresponding SDS-PAGE analysis of the fractions used for experiments, confirming the presence of a predominant protein band at the expected molecular weight for each transporter.

***Supplementary References***

1. Lau, A. M., Claesen, J., Hansen, K. & Politis, A. Deuteros 2.0: peptide-level significance testing of data from hydrogen deuterium exchange mass spectrometry. *Bioinformatics* **37**, 270–272 (2021).

2. Discovery, C. *et al.* Chai-1: Decoding the molecular interactions of life. 2024.10.10.615955 Preprint at https://doi.org/10.1101/2024.10.10.615955 (2024).

3. Hageman, T. S. & Weis, D. D. Reliable Identification of Significant Differences in Differential Hydrogen Exchange-Mass Spectrometry Measurements Using a Hybrid Significance Testing Approach. *Anal. Chem.* **91**, 8008–8016 (2019).
