## Supplemental Tables for "Local conformational plasticity underlies ligand recognition and proton coupling in MFS multidrug transporters"

**Supplementary Table - 1 :** List of peptides from LmrP datasets wild-type (n=3) and mutant D68N (n=1) classified as transmembranous, including TMH number, peptide sequence, start and end positions, and R.F.U. value at 15 min. The last column, "Dynamics", classifies a peptide as flexible when its R.F.U. is above the mean R.F.U. of the extramembranous peptides dataset for at least three biological replicates.

| TMH | Sequence | Start | End | **LmrP WT** R.F.U. (15 min - %) Replicat 1 (Threshold : 29.69%) | **LmrP WT** R.F.U. (15 min - %) Replicat 2 (Threshold : 31.96%) | **LmrP WT** R.F.U. (15 min - %) Replicat 3 (Threshold : 20.07%) | **LmrP D68N** R.F.U. (15 min - %) (Threshold : 29.29%) | Dynamics |
|---|---|---|---|---|---|---|---|---|
| 1 | LRLGIVF | 14 | 20 | / | 21.95% | 6.30% | / | |
| | LRLGIVFL | 14 | 21 | 8.27% | 18.61% | 4.20% | 8.61% | |
| | GIVFLGA | 17 | 23 | 2.48% | 2.58% | 1.97% | 4.9% | |
| | GIVFLGAF | 17 | 24 | 1.39% | / | 2.03% | 2.03% | |
| | GIVFLGAFSYGTVFSS | 17 | 32 | / | 32.94% | 18.85% | / | |
| | GAFSYGTVF | 22 | 30 | 5.43% | 8.14% | 2.53% | 2.69% | |
| | FSYGTVF | 24 | 30 | 8.97% | 10.59% | 3.50% | 5.13% | |
| 2 | GILLALSA | 47 | 54 | 7.99% | / | / | 7.26% | |
| 3 | VFGTIIQL | 78 | 85 | 0.69% | 2.69% | 0.00% | 1.08% | |
| | FGTIIQL | 79 | 85 | 0.06% | / | / | 0.77% | |
| | GTIIQLL | 80 | 86 | 1.15% | / | / | 0.49% | |
| | IQLLGAAL | 83 | 90 | -0.05% | 1.15% | / | 0.16% | |
| | LGAALAI | 86 | 92 | 0.23% | / | 0.65% | 0.36% | |
| 4 | LISFGYN | 111 | 117 | 0.15% | / | / | 0.68% | |
| | ISFGYNF | 112 | 118 | 0.01% | / | / | 2.21% | |
| | FVITAGNA | 118 | 125 | 7.19% | / | 3.92% | 15.73% | |
| | FVITAGNAM | 118 | 126 | 10.87% | 20.41% | 7.91% | 18.76% | |
| | VITAGNAM | 119 | 126 | 14.56% | 26.64% | 10.91% | 23.78% | |
| 5 | SVILGAAL | 150 | 157 | 0.82% | 3.90% | 0.57% | 1.42% | |
| | GAALGAW | 154 | 160 | 8.56% | / | / | 2.10% | |
| 7 | MIFMGAN * | 219 | 225 | 36.22% | 41.86% | 28.19% | 38.25% | Flexible |
| | IFMGAN | 220 | 225 | / | 33.56% | 24.95% | / | |
| | MGANIATT | 222 | 229 | / | / | 36.04% | / | |
| | IIMQFDNF * | 231 | 238 | 36.23% | 35.33% | 25.50% | 29.54% | Flexible |
| 8 | MAIGMIF | 301 | 307 | 11.44% | / | | | |
| 10 | IAGIVY | 319 | 324 | / | 52.42% | / | / | |
| | IAGIVYTLGE * | 319 | 328 | 37.91% | 52.59% | 27.15% | / | Flexible |
| | IVYTLGE * | 322 | 328 | 31.15% | 40.69% | 25.66% | 36.48% | Flexible |
| | IVYTLGEI * | 322 | 329 | 32.45% | 46.88% | / | 33.76% | Flexible |
| | IVYTPSVQ * | 329 | 336 | 35.64% | 42.20% | 29.75% | 35.86% | Flexible |
| | IVYTPSVQTL | 329 | 338 | 42.38% | / | 28.09% | 41.96% | Flexible |
| | IVYTPSVQTLGA | 329 | 340 | 40.74% | / | 28.45% | 40.18% | Flexible |
| | IVYTPSVQTLGAD | 329 | 341 | 36.68% | / | 27.71% | 35.98% | Flexible |
| 11 | AIKMPIASIL * | 356 | 365 | 32.11% | 38.91% | 18.53% | 32.41% | Flexible |
| | IKMPIASIL | 357 | 365 | 7.58% | 36.88% | / | / | |
| | MPIASIL | 359 | 365 | 23.84% | / | 16.98% | 22.49% | |
| | ASILAGL | 362 | 368 | -0.93% | 3.26% | / | 0.79% | |
| 12 | ALTEVLA | 386 | 392 | 9.29% | 29.14% | 1.37% | 10.97% | |

**Supplementary Table - 2 :** List of peptides from NorA WT datasets (n = 3) classified as extramembranous, including TMH number, peptide sequence, start and end positions, and R.F.U. value at 15 min. The last column, "Dynamics", classifies a peptide as flexible when its R.F.U. is above the mean R.F.U. of the extramembranous peptides dataset for three biological replicates..

| TMH | Sequence | Start | End | R.F.U. (15 min - %) Replicat 1 (Threshold : 34.67%) | R.F.U. (15 min - %) Replicat 2 (Threshold : 34.64%) | R.F.U. (15 min - %) Replicat 3 (Threshold : 28.76%) | Dynamics |
|---|---|---|---|---|---|---|---|
| 1 | FLGIGL | 16 | 21 | / | 6.15% | / | |
| | LVIPVL | 21 | 26 | / | / | 14.58% | |
| 2 | LLVAAF | 42 | 47 | / | 2.07% | / | |
| | FALSQM | 47 | 52 | 17.26% | 17.76% | / | |
| | FALSQMIISPFGG | 47 | 59 | 39.58% | / | / | |
| | ALSQMIISPF | 48 | 57 | 50.03% | / | / | |
| | IISPFGGTL | 53 | 61 | 37.69% | 31.59% | 17.35% | |
| 3 | IGLIL | 73 | 77 | 2.78% | / | / | |
| | FSVSE | 78 | 82 | 16.39% | / | / | |
| | FSVSEFM | 78 | 84 | / | 19.76% | 13.25% | |
| 4 | MLSRVIGGM | 95 | 103 | 35.27% | / | / | |
| | MLSRVIGGMSA | 95 | 105 | 37.04% | / | / | |
| | MLSRVIGGMSAG | 95 | 106 | 42.38% | 37.45% | 22.04% | |
| | LSRVIGGMSAG | 96 | 106 | 47.46% | 45.17% | 26.91% | |
| | LSRVIGGMSAGMVMPGVTGL | 96 | 115 | / | / | 26.78% | |
| | SRVIGGMSAG | 97 | 106 | 47.87% | 44.45% | 28.18% | Flexible |
| | SRVIGGMSAGMVMPGVTGL | 97 | 115 | / | 46.98% | 28.09% | |
| | GMVMPGVTGL | 106 | 115 | 51.02% | 48.57% | 40.04% | Flexible |
| | MVMPGVTGL | 107 | 115 | 42.91% | 45.33% | 37.04% | Flexible |
| | VMPGVTGL | 108 | 115 | 48.24% | / | 40.91% | |
| | MPGVTGL | 109 | 115 | / | / | 40.95% | |
| | PGVTGL | 110 | 115 | 61.96% | 58.14% | 47.89% | Flexible |
| 5 | FGYMSA | 129 | 134 | 42.37% | 49.74% | 42.86% | Flexible |
| | GYMSA | 130 | 134 | 36.89% | / | / | |
| | IINSGF | 135 | 140 | 36.73% | 32.47% | 18.59% | |
| | IINSGFILGPGIGGF | 135 | 149 | / | 39.85% | 20.80% | |
| | FILGPGIGGF | 140 | 149 | / | 43.93% | 17.77% | |
| | ILGPGIGGF | 141 | 149 | 43.51% | 40.53% | 28.13% | Flexible |
| | GPGIGGF | 143 | 149 | / | 40.74% | / | |
| 6 | FAGAL | 161 | 165 | 23.64% | / | / | |
| | FAGALGIL | 161 | 168 | / | 18.88% | 18.26% | |
| | FAGALGILA | 161 | 169 | / | / | 17.66% | |
| | GILAF | 166 | 170 | 6.83% | / | / | |
| | AFIMS | 169 | 173 | 7.24% | / | / | |
| 7 | FITPVIL | 204 | 210 | / | 27.57% | 26.25% | |
| | ITPVIL | 205 | 210 | 15.29% | 17.19% | / | |
| | LTLVLSFG | 210 | 217 | / | / | 18.46% | |
| | TLVLSF | 211 | 216 | 11.69% | 6.96% | / | |
| | LVLSF | 212 | 216 | 4.74% | / | / | |
| | LVLSFGLSA | 212 | 220 | / | / | 22.19% | |
| | VLSFGLSA | 213 | 220 | / | / | 23.66% | |
| | LSFGLSAF | 214 | 221 | 46.19% | / | / | |
| | SFGLSA | 215 | 220 | 24.71% | / | / | |
| | SFGLSAF | 215 | 221 | / | 30.23% | / | |
| 8 | IAITGGGIF | 242 | 250 | 43.79% | 32.80% | 16.87% | |
| | ITGGGIF | 244 | 250 | 34.15% | / | / | |
| | GIFGALF | 248 | 254 | / | 3.49% | / | |
| | GIFGALFQ | 248 | 255 | 40.18% | / | / | |
| | GIFGALFQ | 248 | 255 | 40.18% | / | / | |
| 9 | LTFIAWSLLYS | 269 | 279 | 39.05% | / | / | |
| | IAWSLL | 272 | 277 | / | 18.00% | / | |
| 10 | DMIRPAITNY | 307 | 316 | / | / | 35.23% | |
| 11 | NSTFTSMGN | 332 | 340 | 32.14% | / | / | |
| | TSMGNF | 336 | 341 | 39.66% | 34.68% | 23.59% | |
| | TSMGNFIGPLIA | 336 | 347 | / | 36.20% | 24.50% | |
| | TSMGNFIGPLIAGAL | 336 | 350 | / | 45.01% | 27.75% | |
| | IGPLIA | 342 | 347 | 41.48% | 42.37% | 32.86% | Flexible |
| | IGPLIAGAL | 342 | 350 | / | 46.76% | 34.02% | |
| 12 | MAIGVS | 361 | 366 | 21.72% | / | / | |
| | MAIGVSL | 361 | 367 | 21.56% | 36.39% | 46.73% | |
| | AIGVSL | 362 | 367 | 18.76% | 32.67% | 47.30% | |
| | VSLAGVV | 365 | 371 | 15.69% | 25.67% | 33.49% | |
| | AGVVIVL | 368 | 374 | 1.77% | / | 18.04% | |
| | VVIVL | 370 | 374 | 6.43% | / | / | |
| | EKQHRAKL | 376 | 383 | 21.84% | / | / | |

**Suppl. Table - 3 :** List of peptides from QacA WT datasets (n = 3) classified as extramembranous, including TMH number, peptide sequence, start and end positions, and R.F.U. value at 15 min. The last column, "Dynamics", classifies a peptide as flexible when its R.F.U. is above the mean R.F.U. of the extramembranous peptides dataset for three biological replicates.

| TMH | Sequence | Start | End | R.F.U. (15 min - %) Replicat 1 (Threshold : 21.98%) | R.F.U. (15 min - %) Replicat 2 (Threshold : 34.41%) | R.F.U. (15 min - %) Replicat 3 (Threshold : 26.42%) | Dynamics |
|---|---|---|---|---|---|---|---|
| 1 | FVVTMD | 29 | 34 | / | 13.85% | / | |
| | FVVTMDM | 29 | 35 | 17.82% | / | / | |
| | VVTMDM | 30 | 35 | / | 29.33% | / | |
| | IMALPEL | 39 | 45 | 14.73% | 28.65% | / | |
| 2 | WIVDIYS | 58 | 64 | / | 60.60% | 54.93% | |
| | IVDIYS | 59 | 64 | / | 49.61% | / | |
| | AGFIIPLSA | 68 | 76 | 8.72% | 15.13% | / | |
| | FIIPLSA | 70 | 76 | 19.06% | 17.48% | 50.89% | |
| | IIPLSA | 71 | 76 | / | 16.30% | / | |
| 3 | LTGFALFG | 88 | 95 | 22.00% | / | / | |
| | FGLVSL | 94 | 99 | 5.34% | 1.79% | / | |
| 4 | IRFLLGIAGA | 113 | 122 | / | 12.77% | / | |
| | LIMPTT | 123 | 128 | / | 18.06% | / | |
| | LIMPTTL | 123 | 129 | 14.22% | 18.71% | / | |
| | LIMPTTLSM | 123 | 131 | / | 22.23% | 21.52% | |
| | PTTLSM | 126 | 131 | / | 0.75% | / | |
| 5 | AVWSIA | 147 | 152 | / | 20.52% | / | |
| | AVWSIASS | 147 | 154 | / | / | 23.81% | |
| | AVWSIASSIG | 147 | 156 | / | 27.41% | / | |
| | WSIASSIG | 149 | 156 | / | 27.01% | 21.35% | |
| | GAVFGPIIGGAL | 156 | 167 | 7.24% | / | / | |
| | GAVFGPIIGGALL | 156 | 168 | 7.57% | / | / | |
| | AVFGPIIGGAL | 157 | 167 | 4.02% | 22.82% | / | |
| | VFGPIIGGALL | 158 | 168 | 0.51% | / | / | |
| | PIIGGAL | 161 | 167 | / | 16.38% | / | |
| 6 | LINVPFA | 178 | 184 | / | / | 17.95% | |
| | LINVPFAIIA | 178 | 187 | 5.00% | 12.35% | / | |
| | INVPFAII | 179 | 186 | / | / | 12.90% | |
| | AVVAGL | 187 | 192 | 6.00% | 4.50% | / | |
| | AVVAGLF | 187 | 193 | 1.08% | / | / | |
| | VVAGLFL | 188 | 194 | 23.67% | / | / | |
| 7 | SIAGMIG | 216 | 222 | 9.83% | / | 6.38% | |
| | EFSKEGL | 229 | 235 | 53.25% | / | / | |
| 9 | FSAGTIAA | 276 | 283 | 26.75% | / | / | |
| | FAMASVL | 288 | 294 | 7.12% | 9.04% | 13.00% | |
| | FAMASVLL | 288 | 295 | 6.84% | / | / | |
| | LLASQWL | 295 | 301 | / | / | 23.19% | |
| | LASQWL | 296 | 301 | 11.22% | / | / | |
| | ASQWLQV | 297 | 303 | 29.45% | / | / | |
| 10 | LYLLPMA | 314 | 320 | 67.22% | / | / | |
| | YLLPMAIG | 315 | 322 | / | / | 27.39% | |
| | YLLPMAIGD | 315 | 323 | 20.72% | / | 20.44% | |
| | YLLPMAIGDMVF | 315 | 326 | / | / | / | |
| | IGDMVF | 321 | 326 | 31.95% | / | / | |
| | IGDMVFAP | 321 | 328 | 40.31% | / | / | |
| | MVFAPIAPGL | 324 | 333 | / | / | 30.89% | |
| | VFAPIAPGL | 325 | 333 | 20.86% | 29.33% | 32.57% | Flexible |
| | VFAPIAPGLA | 325 | 334 | / | / | 32.12% | |
| | APIAPG | 327 | 332 | 67.61% | / | / | |
| | APIAPGL | 327 | 333 | 29.80% | 37.44% | 36.69% | Flexible |
| | APIAPGLA | 327 | 334 | 37.33% | 36.52% | 39.49% | Flexible |
| 11 | PSGIGIAA | 344 | 351 | 4.47% | 20.28% | / | |
| | AAIGMF | 350 | 355 | 28.28% | / | / | |
| 12 | ILVGAGM | 374 | 380 | / | / | 5.13% | |
| | ILVGAGMASL | 374 | 383 | 8.97% | 7.67% | / | |
| | VGAGMASL | 376 | 383 | / | / | 28.18% | |
| | AVASAL | 384 | 389 | 14.97% | 11.49% | / | |
| | AVASALIM | 384 | 391 | / | / | 15.67% | |
| 13 | PTSKAGNAAA | 395 | 404 | / | 39.10% | / | |
| | YDLGNVF | 410 | 416 | / | 17.17% | 14.27% | |
| | YDLGNVFG | 410 | 417 | 5.41% | 23.00% | 16.74% | |
| | YDLGNVFGVAVL | 410 | 421 | 6.66% | 18.90% | 15.70% | |
| | GSLSSM | 422 | 427 | / | 24.43% | / | |
| | GSLSSML | 422 | 428 | 15.25% | 19.61% | / | |
| 14 | AVTSFNDA | 474 | 481 | 12.97% | / | / | |
| | AVTSFNDAF | 474 | 482 | / | / | 31.97% | |
| | FNDAFVA | 478 | 484 | 38.80% | / | / | |
| | FVATAL | 482 | 487 | / | 11.80% | / | |
| | VATALV | 483 | 488 | 38.80% | / | / | |
| | TALVGGIIM | 485 | 493 | 1.52% | / | / | |
| | ALVGGIIMII | 486 | 495 | / | 10.10% | / | |
| | VGGIIM | 488 | 493 | 1.15% | 1.88% | / | |
| | GGIIMI | 489 | 494 | 42.94% | / | / | |
| | GGIIMII | 489 | 495 | 50.15% | / | / | |
